# CellSP: Module discovery and visualization for subcellular spatial transcriptomics data

**DOI:** 10.1101/2025.01.12.632553

**Authors:** Bhavay Aggarwal, Saurabh Sinha

## Abstract

Spatial transcriptomics has enabled the study of mRNA distributions within cells, a key aspect of cellular function. However, there is a dearth of tools that can identify and interpret functionally relevant spatial patterns of subcellular transcript distribution. To address this, we present CellSP, a computational framework for identifying, visualizing, and characterizing consistent subcellular spatial patterns of mRNA. CellSP introduces the concept of “gene-cell modules”, which are gene sets with coordinated subcellular transcript distributions in many cells. It provides intuitive visualizations of the captured patterns and offers functional insights into each discovered module. We demonstrate that CellSP reliably identifies functionally significant modules across diverse tissues and technologies. We use the tool to discover subcellular spatial phenomena related to myelination, axonogenesis and synapse formation in the mouse brain. We find immune response-related modules that change between kidney cancer and healthy samples, and myelination-related modules specific to mouse models of Alzheimer’s Disease.

## INTRODUCTION

Spatial transcriptomics (ST) technologies and analytical tools available today offer unprecedented views of gene expression patterns within the intricate tapestry of tissues [1] [2] [3]. At the cutting edge of this technology are single-molecule resolution assays [4] [5] [6], which provide a detailed view of transcript distributions within individual cells, promising to transform our understanding of mRNA localization and its relationship with cellular functions. This relationship has been documented in the literature through anecdotal examples showing that RNA localization to specific subcellular regions may underlie efficient spatial organization of cellular processes [7], rapid local translation in response to external stimuli [8], maintenance of cell polarity [9] [10], facilitation of cell migration [11], coordination of developmental patterning [12], etc. Thus, there is growing recognition of the need to expand our understanding of subcellular RNA localization [13], and the time is ripe for this pursuit, supported by the state-of-the-art subcellular spatial transcriptomics tools.

While experimental techniques have advanced rapidly – enabling, for instance the mapping of individual transcripts of hundreds to thousands of genes in thousands of cells [14] [15] [16] – analytical methods have lagged behind. Available ST-related tools are mostly designed for analysis at the single-cell resolution [17]. More recently, methods targeting subcellular phenomena have emerged [13] [18] [19] [20] that can highlight subcellular localization patterns involving single genes or gene pairs, in individual cells. For example, a gene may be annotated as having transcripts localized to cell edges by the BENTO tool [13] or having a “radial” distribution in a cell by the SPRAWL software [19]. Similarly, a gene pair may be found to be significantly colocalized by the InSTAnT toolkit [18], in an individual cell or across many cells. However, when applied to a typical subcellular ST dataset, these tools identify tens of thousands of statistically significant subcellular spatial patterns involving individual genes or gene pairs in individual cells. This overwhelming volume of results imposes a significant interpretive burden, complicating the transition from statistical findings to meaningful and actionable biological insights. Moreover, existing methods are fundamentally limited in scope: they operate at the level of single genes or gene pairs, lacking the ability to capture broader patterns of spatial organization of mRNA. The goal of this work is to bridge this gap with an analytical method that identifies the most salient subcellular spatial patterns in an ST data set, along with biological interpretations of those patterns.

An important lesson from decades of transcriptomics data analysis is the concept of gene “module” – a set of genes that share some common pattern of expression. Systematic identification of gene modules [21] followed by statistical enrichment tests [22] is a popular approach to distil a large number of statistical observations into a more compact set of systems-level insights. Inspired by this paradigm, we sought a method to define gene modules with shared subcellular spatial patterns. One simple strategy is to determine if a gene exhibits a specific subcellular localization (e.g., “cell edge” according to BENTO) in many cells, and group together all genes exhibiting the same frequent localization into a module. This strategy is overly permissive, since two genes annotated with the same frequent localization pattern may not exhibit that common pattern in the same cells. We therefore conceptualized a “gene-cell” module as a set of genes that exhibit the same subcellular pattern in the same cells. However, requiring all genes of a module to exhibit the same subcellular pattern in all module cells (as in [18]) is overly restrictive, and will result in highly fragmented modules. To address this, we introduce a biclustering approach that searches for genes co-exhibiting the same subcellular pattern in a substantial subset of module cells rather than requiring their presence in all module cells, enabling the discovery of more meaningful and coherent gene-cell modules. This algorithm forms the core of the software presented here, called “CellSP”.

CellSP analyzes single-molecule resolution ST data, identifies significant subcellular spatial distribution patterns at the gene level, and distills them into a compact list of gene-cell modules that typically comprise tens of genes and hundreds of cells. It provides specialized techniques for visualizing such modules and their defining spatial patterns. It uses gene set enrichment analysis to describe the genes comprising the module. Additionally, it uses machine learning classification to distinguish module-associated from other cells in the tissue based on their transcriptomic profiles, identifying genes and biological properties (beyond cell type) that characterize module cells.

We used CellSP to analyze ST data sets from four different studies generated using two different technologies. We present the results of these analyses and provide guidance on interpreting the discovered modules. We identify modules associated with myelination in mouse brain tissues across various technologies and find these modules to exhibit differences linked to Alzheimer’s disease. The reported differences are not necessarily detectable through traditional gene expression analyses. We also find modules related to cell-cell adhesion and axonogenesis, and a closer examination of these modules sheds light on interactions among neighboring cells. Use of CellSP on a cancer data set reveals immune-response related gene-cell modules representing subcellular phenomena specific to cancer versus healthy tissue. In summary, CellSP establishes a robust framework for the systematic detection, visualization and comparison of subcellular transcriptomic patterns, coupled with a statistical characterization of their associated biological functions.

## RESULTS

### Overview of CellSP

CellSP is a tool for analysis of spatial transcriptomics (ST) data at single-molecule resolution (**Fig. 1a**). It identifies “gene-cell modules”, where a module is defined as a set of genes exhibiting a specific subcellular spatial distribution pattern (of their transcripts) in a set of cells (We use the terms “gene-cell module” and “module” interchangeably.) Modules are recovered based on statistical criteria and represent the occurrence of interesting, persistent subcellular spatial phenomena in the tissue. There are three main steps involved:

**Fig. 1:**
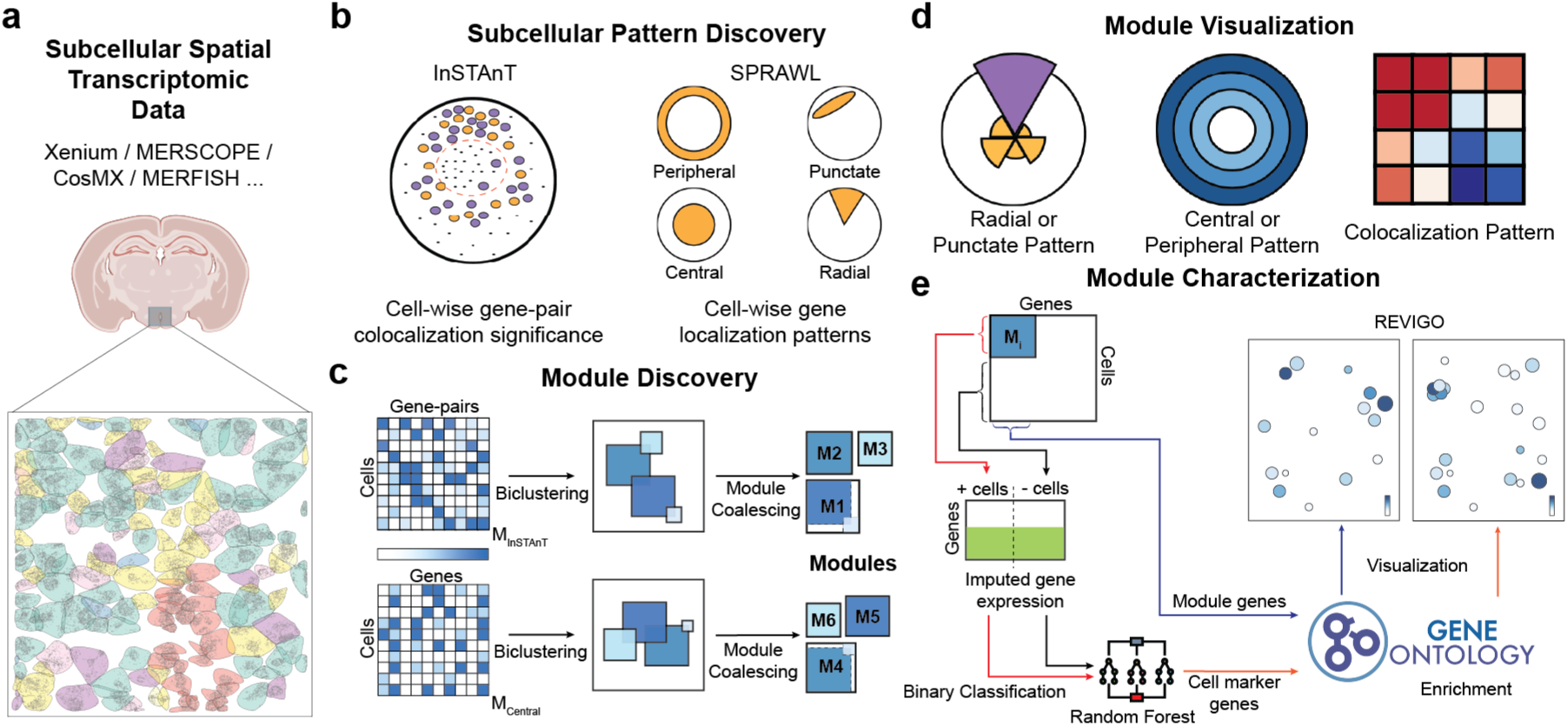
Schematic of CellSP. **(a)** CellSP analyzes single-molecule resolution spatial transcriptomics data from a tissue; such data provides subcellular locations of individual transcripts of a set of genes within each delineated cell. **(b)** Subcellular Pattern Discovery. CellSP utilizes existing tools InSTAnT and SPRAWL for the discovery of subcellular spatial patterns of genes within each cell. Types of patterns detected include gene-gene colocalization reported by InSTAnT and four types of subcellular localization preferences (peripheral, punctate, central, radial) reported by SPRAWL. **(c)** Module Discovery. The next step identifies spatial patterns involving multiple genes and cells. Patterns identified in the previous step (represented as matrices of cells x genes or gene-pairs) are subjected to a biclustering analysis that identifies a subset of rows and columns with a large average value, representing a “gene-cell module”: a set of genes or gene-pairs exhibiting the same subcellular pattern within the same cells. Gene-cell modules that overlap in a significant number of genes (or gene pairs) are coalesced to reduce redundancy. This step outputs a set of gene-cell modules with varying number of cells and genes. **(d)** CellSP provides intuitive visualizations of gene-cell modules, which often span tens to hundreds of cells making direct inspection impractical. CellSP visualizations are tailored to the type of pattern defining the module and summarize the strength of subcellular spatial pattern involving module-associated genes and cells while contrasting them with background genes and cells. **(e)** Module Characterization. Module-associated genes are subjected to Gene Ontology (GO) analysis to gain functional insights into detected gene-cell modules. Alternatively, a Random Forest classifier is trained to differentiate module cells from non-module cells and the most predictive genes, called cell-marker genes, are subjected to GO analysis. To address the limited size of gene panels in subcellular spatial transcriptomic datasets, CellSP employs Tangram to impute gene expression from single-cell RNA-seq datasets of the same tissue, enabling more comprehensive analysis. Results of GO analysis are visualized using REVIGO plots.

*Step 1 – Subcellular pattern discovery*. Here, the statistical tools SPRAWL [19] and InSTAnT [18] are used to rigorously identify subcellular spatial patterns involving individual genes (SPRAWL) or gene pairs (InSTAnT), in each cell (**Fig. 1b**). SPRAWL identifies four types of subcellular patterns – peripheral, radial, punctate and central – describing the distribution of a gene’s transcripts within the cell, while InSTAnT tests if transcripts of a gene pair tend to be proximal to each other more often than expected by chance. Patterns of each type are stored in a matrix whose columns represent genes or gene pairs and rows represent cells (**Fig. 1c**, left), with entries representing strength of pattern occurrence. Five such matrices are produced, one for each of the four spatial patterns from SPRAWL analysis and one from InSTAnT analysis.

*Step 2 – Module discovery*. Next, a statistical tool called LAS (Large Average Submatrices) [23] is used to analyze each pattern annotation matrix and identify “biclusters”, i.e., a subset of rows and columns with a large average value (**Fig. 1c**, middle). Each bicluster represents a set of genes or gene pairs that exhibit the same type of subcellular pattern in the same set of cells, with statistical significance estimated by a Bonferroni-based score (**Methods**). Biclusters identified in this manner may overlap in terms of rows (cells) and columns (genes/gene pairs), yielding many redundant modules. To address this, CellSP deploys an iterative module-coalescing process where pairs of modules comprising similar sets of cells and genes are combined into larger modules (**Methods**, **Fig. 1c**, right). The module discovery step utilizes a parallel computing implementation to accelerate runtime and facilitate the analysis of large datasets.

*Step 3 – Module characterization.* To aid biological interpretation, CellSP reports shared properties of the genes and cells of each discovered module (**Fig. 1e**). Genes are characterized using Gene Ontology (GO) enrichment tests, while cells are characterized by their cell type composition if such information is available. To provide a more precise characterization of a module’s cells, CellSP trains a machine learning classifier to discriminate those cells from all other cells, using the expression levels of all genes other than the module genes. (Here, it uses a reference scRNA-seq data set, if available, to impute levels of genes missing in the ST data set; see **Methods**.) Genes that are highly predictive in this task are then subjected to GO enrichment tests, furnishing hypotheses about biological processes and pathways that are active specifically in the module cells.

#### User experience

The CellSP tool ingests a single-molecule resolution ST data set in the popular “AnnData” format with cell delineations (**Methods**) and returns a listing of gene-cell modules identified for all five pattern types (peripheral, radial, punctate, central, colocalization). The detailed report of each module includes the type of subcellular pattern, identities of genes, number of cells, GO terms characterizing the module genes, cell type composition and GO terms associated with module cells. The latter two are provided depending on the availability of cell type annotations and reference scRNA-seq data (**Methods**), both of which are optional.

#### Visualization

CellSP introduces new techniques for intuitive visualization of subcellular phenomena represented by modules. Here, we outline these visualization methods in general terms, specific examples are provided in subsequent sections. For modules defined by gene pair colocalization, it creates a heatmap showing that colocalization frequencies are higher for module genes versus other genes, and in module cells versus other cells (**Fig. 1d**, right). Modules of genes that localize in “central” or “peripheral” pattern are depicted using a density plot of transcript distribution at varying radial locations in an idealized circular cell, averaged over all module cells (**Fig. 1d**, middle). Subcellular patterns of types “radial” and “punctate” are depicted using a density plot binned by sectors of an idealized circular cell, aggregated over all module cells by aligning their densest sector (**Fig. 1d**, left). In each of these visualizations, a side-by-side visual comparison with non-module genes is provided to underscore the specificity of the pattern to the module genes. Additional visual aids provided in the report include sample cells illustrating the subcellular phenomenon directly, “REVIGO” [24] plots of GO term enrichments, and Uniform Manifold Approximation and Projection (UMAP) [25] as well as spatial plots of module cells.

### CellSP reveals myelination- and axonogenesis-related subcellular spatial phenomena in preoptic area of mouse hypothalamus

In its first demonstrative application, we used CellSP to analyze a MERFISH data set comprising 5,149 cells in the hypothalamic preoptic area (POA) of a mouse brain [26], identifying 38 modules spanning the five pattern types and each module comprising 9 genes and 138 cells on average (**Supplementary Fig. 1a, Supplementary Data 1**). Each gene appears in less than four modules on average (**Supplementary Fig. 1b**) and each cell appears in less than three modules on average (**Supplementary Fig. 1c**), indicating a small degree of overlap among the modules. The modules comprise genes enriched in a range of biological processes and cellular components, especially cytoskeleton and cellular structure regulation, neuronal function and development, and metabolic processes (**Supplementary Data 1**). Note that the underlying subcellular pattern discovery tool InSTAnT reports 358 significant gene pairs (p-value < 1e-3) that colocalize in 35 cells on average, while SPRAWL reports 134 genes having a significant pattern (score threshold = 0.5) in 351 cells on average across the four patterns. The resulting compendium of significant statistical findings reported by these tools, tens of thousands in number, creates a substantial interpretive burden. CellSP effectively distills these patterns into a manageable set of modules, streamlining the crucial step of biological interpretation.

We next illustrate the format in which CellSP reports its discovered modules (**Supplementary Fig. 2**) and how specific biological insights can be gleaned from these. The module “M0_I” consists of the four genes *Sgk1* (serum/glucocorticoid regulated kinase 1), *Ttyh2* (tweety family member 2), *Ermn* (ermin, ERM-like protein) and *Ndrg1* (N-myc downstream regulated 1), and a set of 332 cells (**Fig. 2a**). It was identified using InSTAnT in the pattern discovery step (Step 1, see above), which means that these genes exhibit statistically significant colocalization in the module cells, a selection of which are shown in **Fig. 2b**. Notably, the colocalization is observed in varying subcellular regions such as the nucleus, nuclear periphery, cytoplasm and cell membrane. (See **Supplementary Fig. 1e** for more examples.) **Fig. 2c** illustrates how CellSP helps visualize a colocalization module, highlighting the specificity of the phenomenon to the module’s genes and cells. The module cells are mostly mature oligodendrocytes (67%, **Fig. 2d**) and the module genes are enriched for GO terms such as the molecular function “cytoskeletal protein binding” (p-value = 0.001), the cellular component “myelin sheath” (p-value 0.004) and the biological process “supramolecular fiber organization” (p-value = 0.02) (**Fig. 2e**). (Note that cytoskeletal protein binding and supramolecular fiber organization play important roles in formation and maintenance of myelin sheath [27] [28].) Moreover, according to CellSP’s classification-based characterization (Step 3, see above), the module cells are marked by genes enriched for “ensheathment of neurons” (p-value = E-12) (**Fig. 2f**). Thus, CellSP analysis suggests a connection between subcellular colocalization of the module and the myelination process in mature oligodendrocytes. A possible interpretation is that the mRNAs of these four genes are captured in the process of being transported to the myelin sheath for local translation, as has been recorded for at least two other myelin biogenesis-related genes, viz., Mbp (Myelin Basic Protein) [29] and Mobp (Myelin Oligodendrocyte Basic Protein) [30]. Interestingly, another CellSP-reported module – “M11_S” – represents a punctate subcellular pattern (**Supplementary Fig. 1d**) involving three of the four above genes, viz., *Sgk1*, *Ttyh2*, *Ermn*, and another myelination-related gene, *Gjc3* (gap junction protein gamma 3) that has been shown to localize in myelin sheaths [31].

**Fig. 2:**
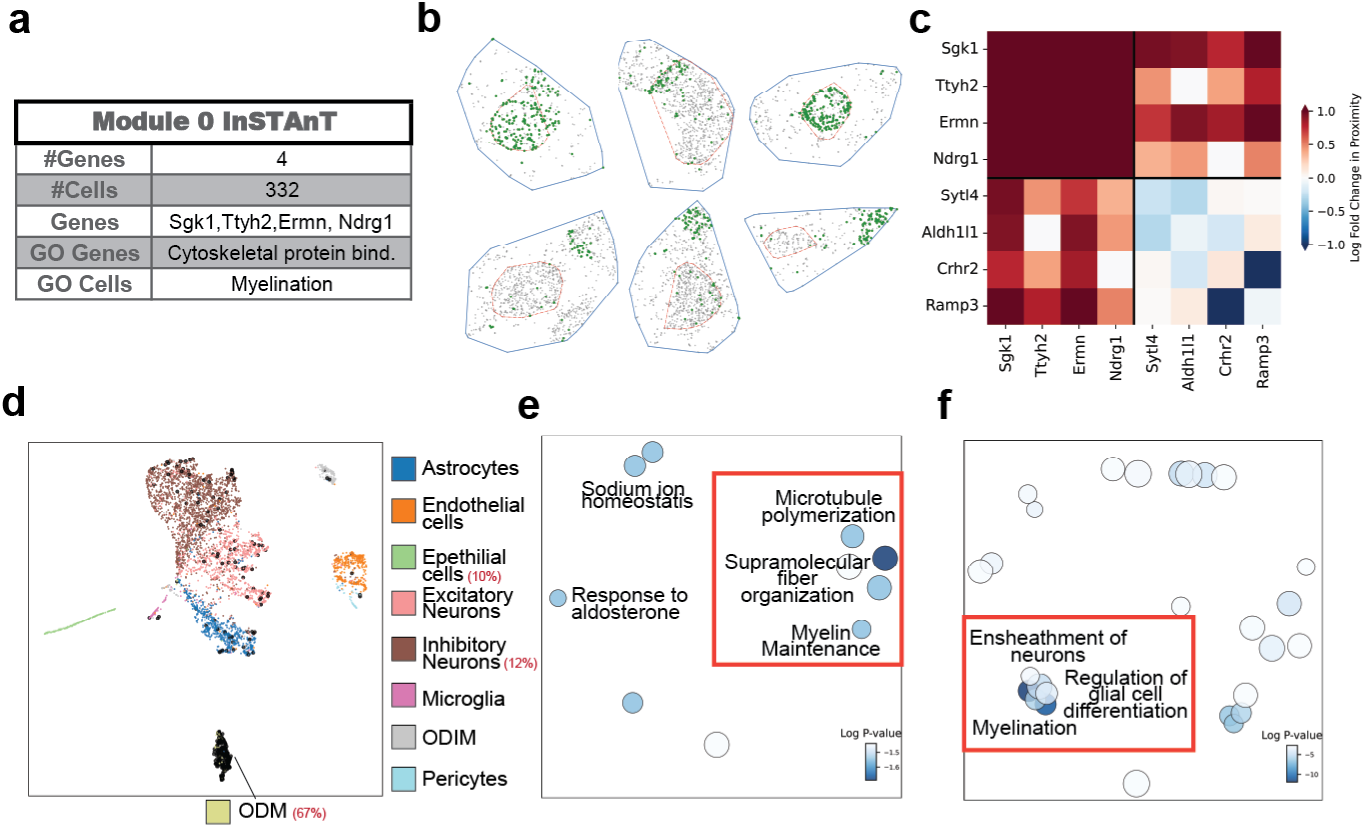
CellSP reveals myelination-related subcellular spatial phenomena in mouse brain. **(a)** Summary table of an example gene-cell module, “M0_I”, discovered by CellSP from MERFISH data on hypothalamic preoptic area in mouse brain. (The suffix “_I” indicates that it was discovered by aggregating single-cell level colocalization patterns reported by InSTAnT.) “GO Genes” shows the Gene Ontology (GO)-based characterization of the four module genes. “GO Cells” shows the GO-based characterization of the genes whose expression levels are most discriminative of the 332 module cells versus remaining cells. **(b)** Direct visualization of six of the 332 module cells, with blue boundaries indicating automatically delineated cell periphery, red boundaries indicating nuclear periphery, each dot representing a transcript, colored in green for module genes and in gray for other genes. **(c)** CellSP visualization of module M0_I. The heatmap shows pairwise colocalization propensity of gene pairs in module cells compared to non-module cells, as a log ratio, with warmer colors (more positive values) indicating specificity of the colocalization to the module cells. The first half of rows and columns corresponds to module genes while the second half corresponds to a random subset of non-module genes, so the contrast between the upper left quadrant (more warm color cells) and other quadrants underscores that the phenomenon is specific to the module gene pairs. **(d)** UMAP visualization of all cells in the sample, with colors indicating cell types and module cells shown in black. The module mostly comprises mature oligodendrocytes (ODM). **(e)** REVIGO plot of biological processes enriched in module genes, showing a prominent association with myelination-related terms. **(f)** REVIGO plot of biological processes associated with gene markers of module cells, showing that these cells are also characterized by genes enriched for myelination-related terms.

An example of a module discovered using SPRAWL in the pattern annotation step is “M0_S” (**Fig. 3a**), consisting of four genes – *Fn1* (fibronectin 1), *Slco1a4* (solute carrier organic anion transporter family member 1a4), *Rgs5* (regulator of G protein signaling 5), *Sema3c* (semaphorin 3C) – distributed in a peripheral pattern (**Fig. 3b**) in 221 cells. **Fig. 3c** illustrates how CellSP helps visualize such a module, revealing the greater tendency of this module’s genes to have peripheral localization, compared with background genes. The module genes are enriched in regulation of axonogenesis (**Fig. 3e**) [32] [33] [34], while the module cells, mostly neurons (50% inhibitory and 30% excitatory, **Fig. 3d**) are marked by genes associated with cell adhesion (**Fig. 3f**), a key process during axonogenesis. The peripheral localization pattern defining the module suggests local translation of proteins that localize in neurites (cell delineations mostly capture somata [35]) or extracellular matrix (ECM). Consistent with this speculation, RGS5 protein is known to exhibit strong synaptic localization [36] [37], while FN1 protein is an extracellular matrix component [38], supporting the possibility of local translation. Considering the module’s statistical association with cell-adhesion functions, we examined the transcript distribution in adjacent pairs of cells and observed that the peripheral localization tends to be at the facing boundaries (**Fig. 3g-i**, **Supplementary Fig. 3**), a phenomenon not seen in non-module cells. Putting together the different facets of this module’s characterization by CellSP, we hypothesize that the four genes are locally translated into proteins that localize at soma boundaries and perform adhesion-related functions. This may be the case if the module cells, mostly neurons, are actively forming synapses, undergoing neurite outgrowth, etc. [39], which is plausible considering that the preoptic region profiled through these data is an important site of developing neurites [40].

**Fig. 3:**
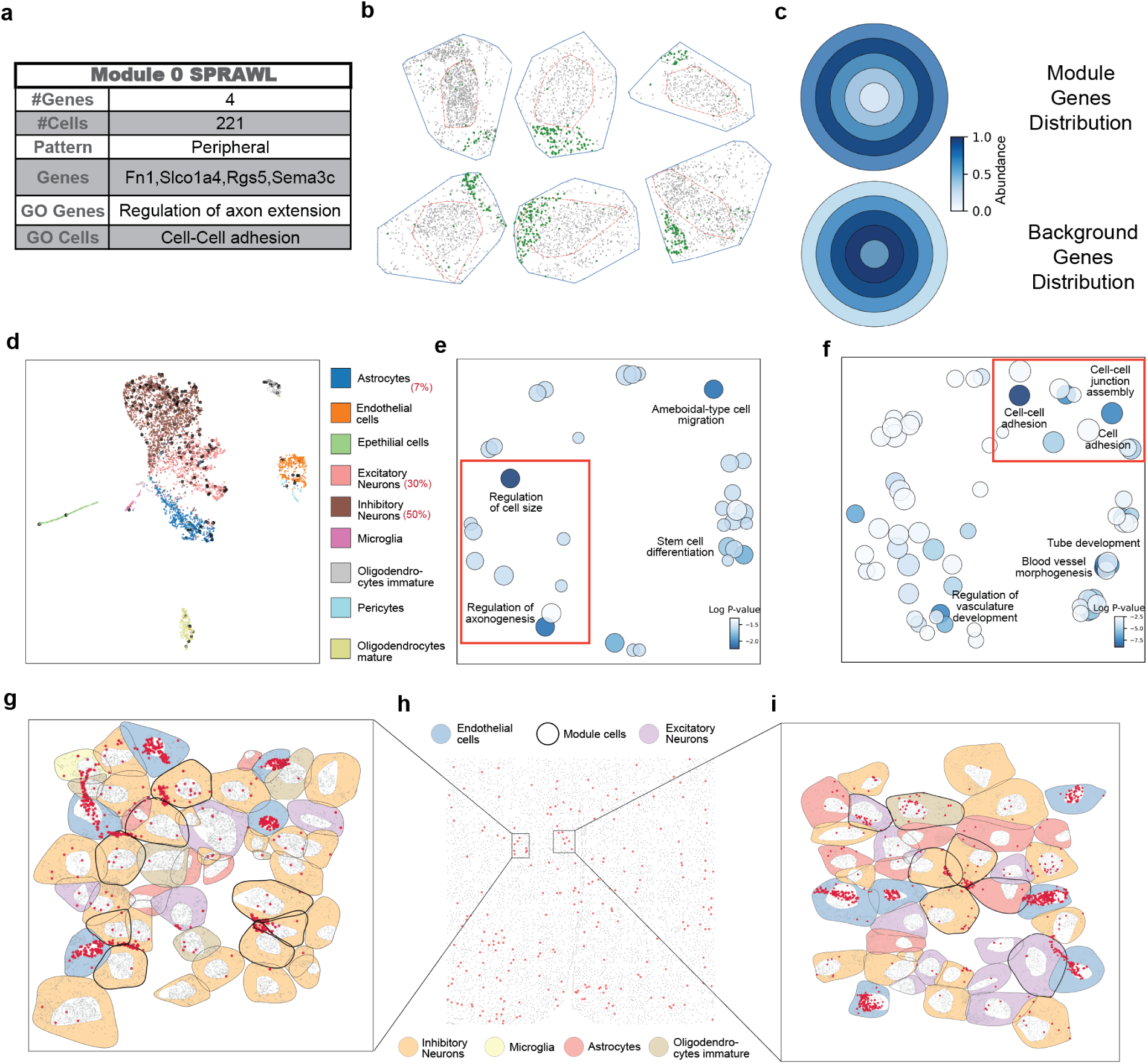
CellSP reveals axonogenesis related subcellular spatial phenomena in mouse brain. **(a)** Summary table of an example gene-cell module, “M0_S”, discovered by CellSP from MERFISH data on hypothalamic preoptic area in mouse brain. (The suffix “_S” indicates that it was discovered by aggregating single-cell level colocalization patterns reported by SPRAWL.) **(b)** Direct visualization of six of the 221 module cells, with each dot representing a transcript, colored in green for module genes and in gray for other genes. **(c)** CellSP visualization of module M0_S. The subcellular space is idealized as a circle with concentric rings representing varying distances from the “center” (see Methods) and color intensity representing transcript abundance of a gene set in a ring, averaged over the module cells. The upper circle depicts this information for module genes and the lower circle represents all other genes for contrast, which in this case highlights that the module gene transcripts are enriched in peripheral regions (outermost ring). This type of visualization is used for any gene-cell module with peripheral or central pattern. **(d)** UMAP visualization of all cells in the tissue, with colors indicating cell types and module cells shown in black. The module mostly comprises neurons. **(e)** REVIGO plot of biological processes associated with module genes, highlighting enrichment for axonogenesis-related terms. **(f)** REVIGO plot of biological processes associated with marker genes of module cells, showing that these cells are characterized by genes enriched for cell adhesion-related terms. **(g-h)** Spatial plot of all cells in tissue (h) with module cells highlighted in red, along with detailed visualization of a subset of module cells in close spatial proximity (g,i). In these detailed views, module cells are depicted with a stronger boundary, while their fill color corresponds to their cell type. Transcripts of module genes are highlighted in red and are localized at the contact boundaries between module cells.

### Subcellular spatial patterns recur across cell types and regions of brain

We next analyzed a large ST data set that profiles a whole mouse brain using Xenium technology [41]. This profiles a panel of 248 genes in 162,033 cells grouped into 50 clusters based on their transcriptomes (**Supplementary Fig. 4**). We used CellSP to analyze each cell cluster separately, as large expression differences across clusters may confound the discovery of subcellular pattern modules, and because we sought to test module reproducibility across varying cellular contexts. CellSP identified ∼22 modules per cell cluster, each module comprising 8 genes and 68 cells on average (**Supplementary Data 2**). Some of the prominent biological characterizations (GO terms) reported for these modules included synaptic processes, myelination, hormone receptor activity, cellular responses to external stimuli, extracellular components, secretory processes and inter-cellular communication.

Closer inspection of these results revealed that modules with certain biological characterizations recur in multiple cell clusters (cell types and/or regions) of the brain. For instance, we observed 36 modules, across 18 different clusters (**Supplementary Data 3, Supplementary Fig. 6**) to be associated with myelination process, i.e., CellSP reported the GO term “ensheathment of neurons” as enriched (nominal p-value < 0.01) in module genes as well as in marker genes of module cells, for each of these modules. There is substantial overlap in their constituent genes (**Fig. 4C**), with the three most frequently recurring constituents being the myelin-associated genes *Gjc3*, *Sox10* (SRY-box transcription factor 10) and *Opalin* (oligodendrocytic myelin paranodal and inner loop protein) [42] [43] [44]. *Gjc3* is known to be expressed in myelinating glial cells [31], *Sox10* is a transcription factor responsible for the activation of several myelin-specific genes [45] and *Opalin* is known to promote oligodendrocyte differentiation and axon myelination [46]. We confirmed that the inclusion of a gene in a module is not a mere reflection of its expression levels in module cells (**Supplementary Fig. 5a**), though higher expression does play a role (**Supplementary Fig. 5b**), in that a gene that is less expressed in a cell is less likely to have its subcellular pattern, if any, detected by a statistical procedure. These myelination-associated modules comprise 3493 cells overall, which are observed across clusters with diverse transcriptomic profiles (**Fig. 4a**) and in different brain regions (**Fig. 4b**). The shared biological properties of these modules are further elucidated by the numerous other GO terms enrichments common to them (**Fig. 4d,e**), including mesenchymal stem cell (MSC) differentiation and oligodendrocyte differentiation. MSCs have the potential to differentiate to or stimulate the maturation of oligodendrocytes [47], the primary myelinating cells.

**Fig. 4:**
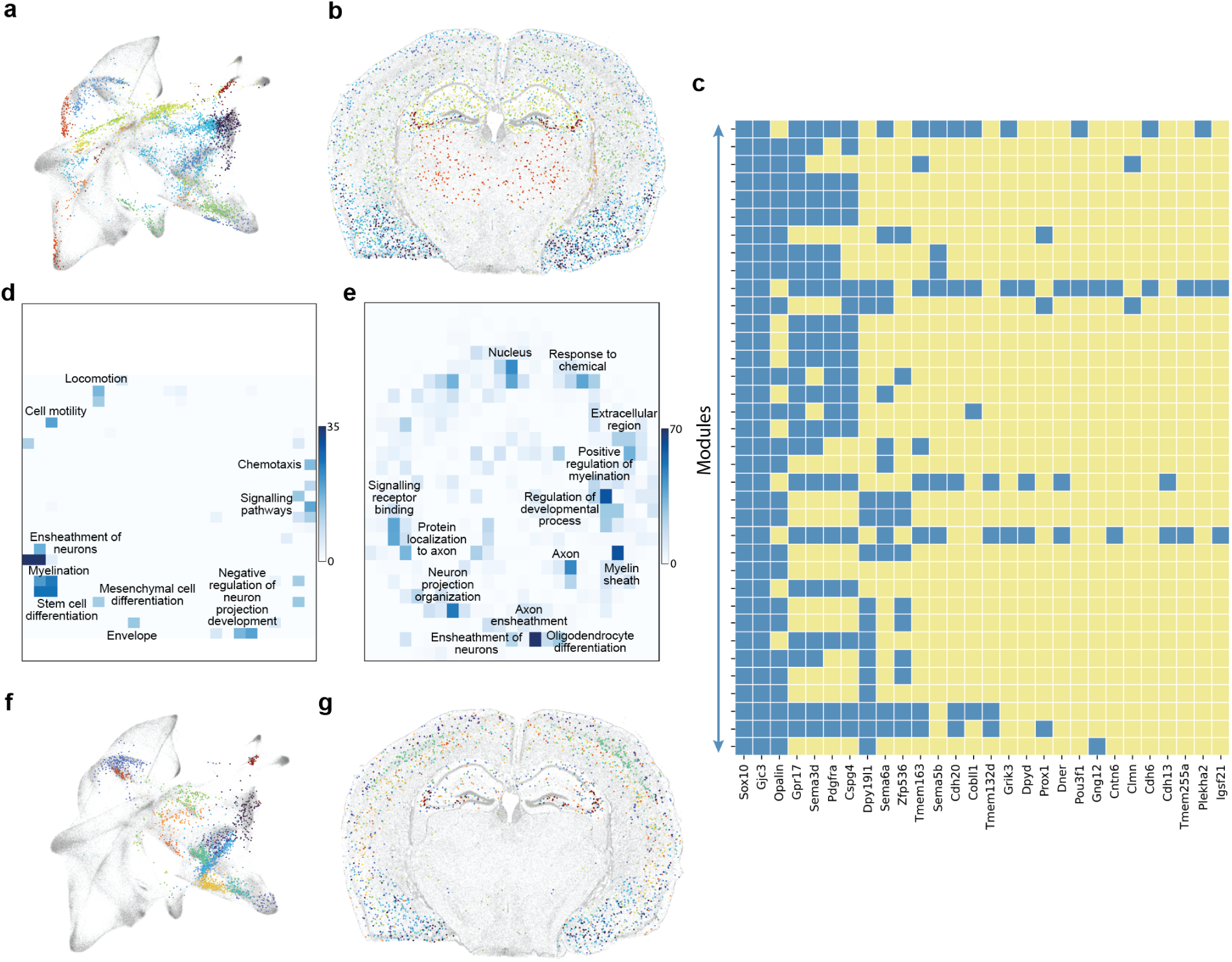
CellSP uncovers myelination and GABA-ergic synapse related modules across cell types and regions of mouse brain. **(a)** UMAP visualization of all cells in a Xenium data set of whole mouse brain, with cells of 36 myelination-related gene-cell modules detected by CellSP shown in color. (Cells of each module are shown in a different color.) This demonstrates that CellSP captures myelination-related subcellular patterns across cells with diverse gene expression profiles, indicating that these patterns are not merely a reflection of overall gene expression levels. **(b)** Spatial plot of all cells in the whole brain data set, with cells of myelination-related modules (same as in A) shown in different colors, while cells not in any of these modules are shown in grey. **(c)** Matrix showing gene composition of each of the 36 myelination-related modules (rows); blue indicates a gene’s inclusion in a module. Genes Sox10, Gjc3, and Opalin appear in all or most modules, while four others appear in a majority of modules. **(d)** REVIGO plot of biological processes associated with module genes, aggregated over all 36 modules. Color intensity of a bin in this plot represents the number of significant module associations of one or more GO terms that map to that bin in the semantic space computed by REVIGO. **(e)** REVIGO plot of biological processes associated with gene markers of module cells, aggregated over all the 36 modules. **(f)** UMAP visualization of all cells in data set, with cells in any of the 20 discovered GABA-ergic synapse-related modules shown in color. (Cells of each module are shown in a different color.) **(g)** Spatial plot of all cells in the sample, with cells from all 20 GABA-ergic synapse-related modules shown in different colors, while cells not in any of these modules are shown in grey.

Another prominent theme in the whole brain CellSP results was the repeated finding of modules characterized by “GABA-ergic synapse”, a GO subcellular localization term. Twenty modules across 14 different cell clusters were reported to have this annotation enriched (p-value < 0.0001) in their constituent genes as well as in expression markers of their member cells (**Supplementary Fig. 7a,b**). These modules were observed in different expression contexts (**Fig. 4f**) and physical regions of the brain (**Fig. 4g**), and to exhibit different types of subcellular patterns (**Supplementary Fig. 9, Supplementary Data 6**). Several genes appear repeatedly as members of these modules, with the most frequent ones being *Gad1* (glutamate decarboxylase 1), *Gad2* (glutamate decarboxylase 2) and *Rab3b* (RAB3B, member RAS oncogene family) (**Supplementary Fig. 7c, Supplementary Fig. 8**), which play critical roles in maintaining basal levels of GABA (Gamma-aminobutyric acid), as well as synthesis and vesicle release of GABA in an activity-dependent manner at inhibitory synapses [48] [49] [50]. There is evidence suggesting that GAD2 and RAB3B proteins are localized in presynaptic terminals [49] [51], hinting at an explanation for the subcellular colocalization of their transcripts.

### CellSP-detected modules can be specific to biological conditions

The analyses above show us that gene-cell modules recovered by CellSP reveal subcellular transcriptomic phenomena related to biological processes such as myelination and axonogenesis. This raises the natural question: do these modules also reveal subcellular phenomena that vary under different tissue conditions? To investigate this, we used CellSP on data sets comprising case-control pairs of tissues. The first such analysis involved Xenium data on a kidney cancer (papillary renal cell carcinoma) FFPE sample (**Supplementary Fig. 10a**) and a healthy kidney sample [52]. As in the whole-brain analysis above, we first clustered the cells of each sample based on their transcriptomes and analyzed each cell cluster with CellSP, identifying 11 modules in the cancer sample and 15 modules in the healthy sample (**Supplementary Data 5, S6**). The cancer modules were mostly (10/11) found in two of the 18 cell clusters (**Fig. 5a, Supplementary Fig. 10b**) – Cluster 6 and Cluster 9. Six of these cancer modules comprise genes and cells characterized by CellSP as being immune response-related, with enrichment of GO terms such as “immune response”, “defense response”, “T cell activation”, “positive regulation of leukocyte proliferation”, etc. (**Supplementary Data 5**).

**Fig. 5:**
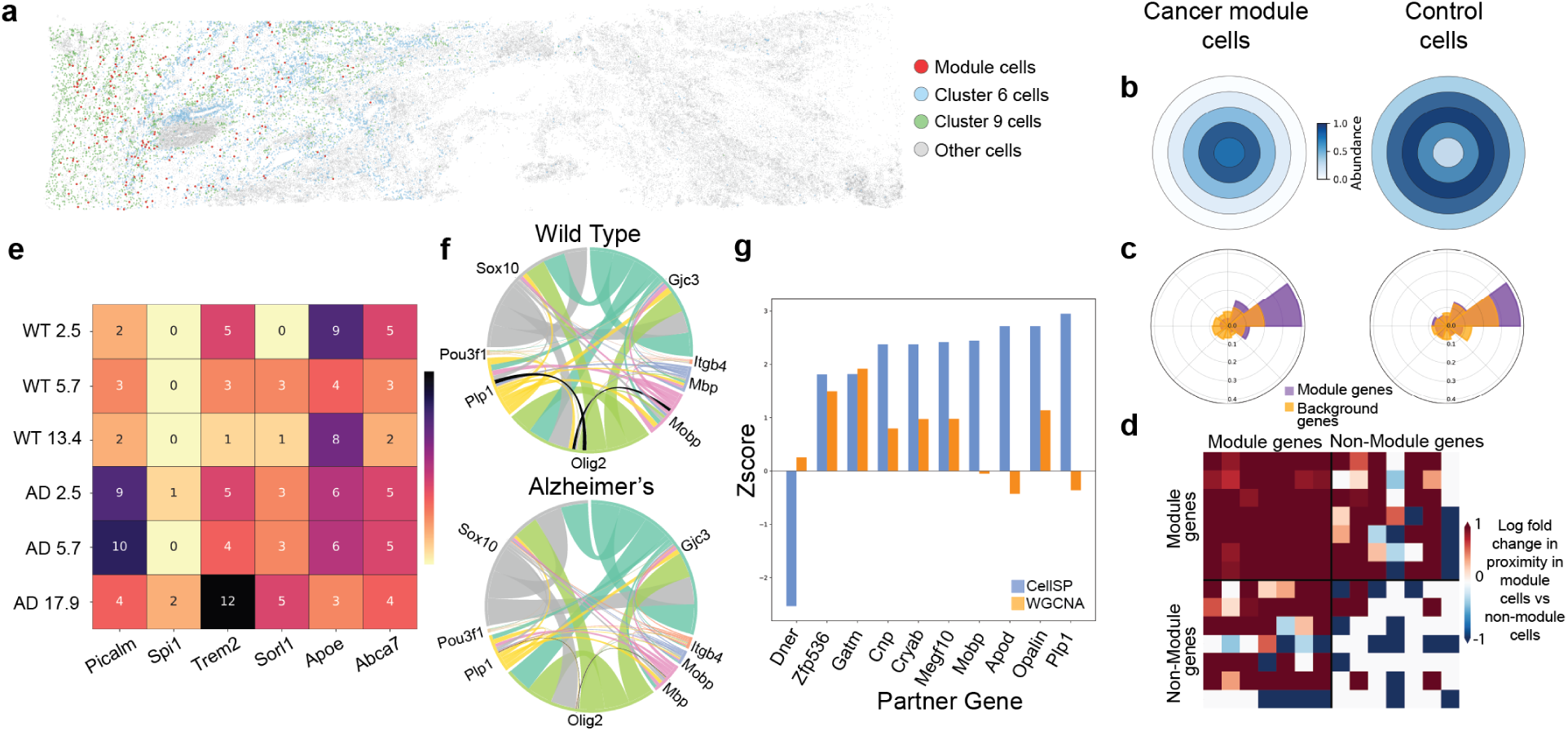
CellSP detects differences in subcellular spatial patterns between tissue conditions. **(a)** Spatial plot of a Kidney PRCC (kidney cancer) tissue (Xenium data), highlighting cluster 6 cells in blue, cluster 9 cells in green, and cells of any of the 6 CellSP-detected immune-response related gene-cell modules in red. **(b, c, d)** CellSP visualization of three selected immune system related gene-cell modules discovered in the cancer sample. (b) shows module R6_M5_S (central pattern) in left panel while right panel is a visualization of the subcellular distribution of the module genes in expression-matched cells from the healthy kidney sample (“control”). Module genes are more uniformly localized towards the cell’s “center” in the cancer sample than in healthy sample. (c) shows module R6_M0_S (radial pattern) in left panel and the right panel depicts the subcellular distribution of module genes in cells from the healthy sample. The cancer sample (left panel) shows a strong clear difference between module genes and non-module genes in terms of radial spread of transcripts, while this contrast is insignificant in the healthy sample (right panel). (d) shows module R6_M3_I, defined by a colocalization pattern. The colocalization visualization has been modified to provide contrast with control cells rather than non-module cancer cells. The module genes are much more proximal to each other in cancer module cells than in random cells from the healthy sample. **(e)** Gene-cell modules detected in mouse hemibrain Xenium samples from three TgCRND8 mice (AD model) at different time points and three wildtype (WT) mice at similar time points were examined for inclusion of six AD-related genes (columns). Shown are the number of detected modules in each of the six samples (rows) that include a gene. Notably, Picalm and Trem2 have more modules detected in AD samples than in WT, consistent with their known roles in Alzheimer’s Disease-related processes. **(f)** Chord diagram depicting the number of modules detected in AD samples (bottom) or WT samples (top) that include a pair of genes, shown for genes annotated with function “ensheathment of neurons”. The gene pairs “Olig2-Plp1” and “Olig2-Mobp” (shown in black) exhibit a significant difference between AD and WT samples. **(g)** Blue bars show z-scores computed from a proportion test comparing AD and WT samples in terms of the fraction of CellSP modules that include Olig2 and another gene (x-axis labels), shown only for partner genes where the proportion test yields a significant p-value. Orange bars show similarly calculated z-scores for modules defined by cell-level co-expression, as computed by WGCNA.

Examination of the 15 modules in the healthy sample (**Supplementary Data 6**) suggested that the immune response-related modules found in the cancer sample were specific to it and were not present in the healthy sample. To pursue this observation objectively, we used CellSP visualization routines to contrast a module’s spatial pattern in the module cells (from the cancer sample) versus cells in the other (healthy) tissue sample. As condition-specific gene expression can be a confounder for such analysis, this analysis used a subset of healthy tissue cells that matched the cancer module cells in the expression levels of module genes (**Methods**). **Figures 5b-d** show the cancer-versus-healthy tissue comparison for three of the seven cancer-associated immune-response modules. For instance, module C6_M5_S (**Fig. 5b**) comprises a set of 31 genes that are localized centrally in a subset of 22 cells (**Supplementary Fig. 10c**) and was detected in Cluster 6 of the cancer sample (**Fig. 5a**). The genes are enriched for the GO term “defense response” (p-value 8.9E-08). As shown in **Fig. 5b** (right), the same set of genes lacks this centrally localized distribution of transcripts in expression-matched cells from the healthy sample. A second example is the module C6_M0_S (**Fig. 5c**), a set of 24 genes, also enriched in “defense response” (p-value 3.3E-04), that exhibit radial subcellular localization in a set of 18 cells (**Supplementary Fig. 10d**) of Cluster 6. CellSP visualization (**Fig. 5c**, left) confirms that the spatial pattern is significantly different between module genes and other genes when examining the module cells. The adjacent panel (**Fig. 5c**, right) shows that this prominent contrast is not seen in expression-matched cells from healthy tissue, supporting the cancer-specificity of this module. The third cancer-specific module we highlight here is C6_M3_I (**Fig. 5d**), a set of seven genes enriched in the term “T cell activation”, that exhibit significant subcellular colocalization in 66 cells (**Supplementary Fig. 10e**) of Cluster 6. The strength of its spatial pattern in cancer versus healthy tissue is visualized in **Fig. 5d**, where the top left quadrant of the heatmap indicates that the module genes have a stronger tendency for pairwise colocalization in the module (cancer) cells compared to expression-matched non-module cells from the healthy sample. (The other quadrants, especially the bottom-right, show that this contrast is not observed for a random subset of non-module genes.) In summary, CellSP discovers immune-response related gene-cell modules representing significant subcellular patterns that are specific to cancer versus healthy tissue.

### CellSP detects subcellular patterns associated with Alzheimer’s Disease in a mouse model

We next used CellSP to identify gene-cell modules in brains of mouse models of Alzheimer’s Disease (AD). For this, we analyzed Xenium data on a coronal section of one hemisphere of TgCRND8 transgenic male mouse brain at three time points (pathological progression stages), as well as wild-type (WT) mouse brain at similar time points [53]. Following the same procedure as above, we clustered cells in each brain sample and used CellSP to find significant modules in each cell cluster, recovering ∼11 gene-cell modules per cluster on average (**Supplementary Fig. 11**, **Supplementary Data 7**). To contextualize these modules with the AD genetics literature, we focused on six genes from the Xenium gene panel implicated in AD risk [54] (**Fig. 5e**). We observed the gene *Picalm* (phosphatidylinositol binding clathrin assembly protein) to appear in many modules in the AD mice, especially in the early and middle time points, significantly more frequently than in WT mice (t-test p-value 0.047) (**Supplementary Fig. 12**). The Picalm protein is responsible for recruiting clathrin and adaptor protein-2 (AP-2) to the plasma membrane [55] as part of clathrin-mediated endocytosis, and contributes to clearance of amyloid-β (Aβ) at the blood brain barrier [56]. It is linked to processes that are disrupted in AD and is also genetically associated with the disease [57]. We thus speculate that the observed subcellular patterns involving this gene are reflections of the gene’s dysfunction in AD mice. We observed another AD risk gene – *Trem2* (triggering receptor expressed on myeloid cells 2) – to feature in many CellSP modules (12) (**Supplementary Fig. 13**) in the late-stage AD mouse brain, significantly more than in the other five samples from WT or AD mice. This late stage is associated with increase in AD-associated microglial population [58], and *Trem2* is primarily expressed in microglia, where it functions as a transmembrane receptor [59] enabling the progression to a mature disease-associated microglia phenotype [58] and is involved in the response to Aβ plaques [60]. Variants in *Trem2* gene have been identified as a significant risk factor for late-onset AD [61], suggesting again that the observed gene-cell modules reveal subcellular phenomena reflecting its dysfunction. The four other AD risk genes were found in few modules overall (**Fig. 5e**) and these modules were of similar counts in AD vs. WT mice. We also note that the above AD-associated changes involving *Picalm* and *Trem2* expression, observed at the subcellular spatial level, would not have been detected at the level of overall cellular expression (**Supplementary Fig. 14**).

We next examined the CellSP-detected modules in AD and WT brains with a focus on myelination. Given the reported connection between myelin damage and AD pathology [62]; and building on our earlier identification of myelination-related gene-cell modules in the mouse brain, we investigated whether such modules have any unique characteristics in one phenotypic group compared to the other. We concentrated on all identified modules associated with the GO term “ensheathment of neurons” (nominal p-value < 0.01) and, to identify any characteristics that differentiate these modules between AD and WT, we examined the frequency of myelination-related genes co-occurring in the same module (**Fig. 5f**). While most gene pairs show similar co-occurrence statistics between the two groups, the pairs *Olig2* (oligodendrocyte transcription factor 2)-*Plp1* (proteolipid protein 1) and *Olig2*-*Mobp* have a significantly higher co-occurrence in WT brain compared to AD brain (proportion test p-value < 0.05). (Also see **Supplementary Data 8**). To follow up on these intriguing observations involving *Olig2*, a gene essential to maturation of oligodendrocytes (the cells responsible for producing myelin), we next identified all genes that co-occur with *Olig2* in a group-specific manner (proportion test p-value < 0.05). (The previous analysis was limited to eight select myelination-related genes, including *Olig2*.) We found 10 such genes (**Fig. 5g**, blue bars), nine of which co-occur with *Olig2* preferentially in gene-cell modules of the WT brains, and consist of the above-noted genes *Plp1*, *Mobp* as well as other genes linked to myelination, including *Opalin* and *Cnp* (2’,3’-cyclic nucleotide 3’ phosphodiesterase) [46] [63] [64] [65]. This inter-group difference in module composition is most in time points 1 and 2 (**Supplementary Data 9**) which correspond to periods of continued myelination in the mice brain [66]. These observations suggest that the integrity and functionality of subcellular modules found in WT brains may be impaired in AD brains, reflecting a broader dysfunction in myelination processes in AD [67]. In summary, our myelination-focused examination of CellSP modules points to a potential significance of *Olig2* in Alzheimer’s disease pathology, highlighting a shift from normal myelination functions in healthy brains.

The specific relationships and insights uncovered here are a glimpse into the dynamics of the subcellular transcriptome during AD progression, not merely a reflection of differential co-expression of genes at the whole cell level. To substantiate this, we repeated the module discovery using co-expression analysis with the popular WGCNA tool [21]. Modules were detected for each cell cluster in each of the six brain samples, as above, except that these were gene co-expression modules rather than CellSP-identified gene-cell modules. These co-expression modules do not show any preferential co-occurrence of *Olig2* with any of genes identified above except one – *Olig2*-*Gatm* (glycine amidinotransferase) (p-value < 0.05, **Fig. 5g**, orange bars). This analysis underscores the ability of CellSP modules to reveal subcellular pattern changes between phenotypic conditions, that may not be discernible at the cellular level.

## DISCUSSION

With the increasing popularity of spatial transcriptomics (ST) techniques of single-molecule resolution [68] [69] [70] new tools have been proposed to identify genes and gene pairs that exhibit interesting transcript distribution patterns within cells [18] [19] [13]. However, there remain fundamental and practical challenges not addressed by these analysis tools. A practical problem is that they annotate subcellular patterns such as gene-gene colocalization or gene localization preferences in individual cells, leading to very large compendia of significant patterns that are difficult to sift through for actionable insights. A conceptual limitation is that while they can effectively shortlist genes that merit further exploration on account of their subcellular distributions, they do not highlight subsets of cells where interesting spatial phenomena manifest and could point to special functional properties of those cells. The simple approach of examining every cell that exhibits a statistically significant spatial pattern is impractical, as a very large number of cells harboring diverse patterns of varying functional origins are enumerated in this way. On the other hand, examining all cells where a particular gene (or pair) has the same subcellular distribution pattern will demand that we repeat such examination for a large number of genes, again leading to a substantial interpretive burden.

The key innovation of CellSP is the idea of a “gene-cell module”, which is one solution to the above shortcomings of existing tools. Such a module, if found to be statistically significant, draws our attention to a subset of cells that share a subcellular spatial pattern involving the same subset of genes, suggesting a functional commonality among those cells. Furthermore, CellSP automatically searches for that functional characterization by using machine learning to identify gene markers of the member cells of the module, and reporting biological properties enriched in the marker genes. This is how our analysis of MERFISH data on hypothalamic preoptic area uncovered a collection of 332 cells, mostly mature oligodendrocytes, that appear to be involved in myelination, and another subset of 221 cells, mostly neurons, that are characterized by axonogenesis-related functions. These discoveries would not have been possible with existing tools. In identifying a gene-cell module, CellSP not only highlights an interesting subset of cells, it simultaneously draws our attention to a set of genes that share an interesting transcript distribution pattern in those cells, prompting functional hypotheses involving those spatial patterns. Such gene set discovery is a standard technique of transcriptomics analysis, e.g., as co-expression module identification [21] and a CellSP module extends this time-tested concept to subcellular colocalization of genes. It was this functionality that led to the identification of the four-gene module comprising myelination-related genes (*Sgk1*, *Ttyh2*, *Ermn* and *Ndrg1*) that colocalize with each other in many mature oligodendrocytes, and another module comprising axonogenesis-related genes (*Fn1*, *Slco1a4*, *Rgs5*, *Sema3c*) whose transcripts localize at the periphery in the same collection of cells, raising the possibility of localized translation of these genes as part of cell adhesion processes.

We note that the InSTAnT toolkit [18] provides module discovery functionality, where a module has a different statistical interpretation from that adopted in CellSP – it is a subset of genes that colocalize with each other significantly frequently across the entire population of cells. We compared CellSP to the gene module discovery methods implemented in InSTAnT (**Supplementary Note 1**) and found that the former offers significant advantages in terms of biological interpretation and ease of use. Importantly, CellSP-reported modules include four additional types of subcellular spatial patterns (radial, peripheral, punctate and central patterns discovered by SPRAWL [19]) and are accompanied by innovative visualizations and machine learning-based characterization of module cells, features that are crucial to the scientific discovery process involving functional subcellular patterns.

We recognize that certain aspects of CellSP can be further improved in future versions. For example, the heuristic nature of the LAS search algorithm and our module merging step may result in variability across different runs, and alternative search algorithms should be explored to address this. As another example, the scheme for visualizing subcellular idealize cells as unit circles, which may not accurately depict elongated or irregularly shaped cells; techniques for average shape construction [71] may be fruitful for future work in this direction. Finally, the comparison of gene-cell modules between conditions is not implemented as a statistical tool in CellSP and was performed by us through downstream processing of its results; such a tool for “differential module discovery” will be another important goal of future work.

By analyzing data sets from diverse organs and tissues, such as whole brain, specific brain regions, kidney, etc., under various biological conditions such as Alzheimer’s disease models or kidney cancer, and different technologies such as MERFISH and Xenium, we demonstrated how CellSP can be used to explore large and high-dimensional ST data and extract new insights into subcellular transcript distribution. The discovered patterns are suggestive of co-transportation or co-localization of RNAs that participate in similar biological functions [72], and also support the phenomenon of local translation [73] which requires the mRNA to be transported to precise locations within the cell where they are translated into proteins. Furthermore, we showed how gene-cell modules can be compared across conditions to discover subcellular changes associated with disease states, which are not recover-able using traditional differential expression analysis. Overall, CellSP offers a powerful new approach to subcellular spatial transcriptomics data analysis.

## METHODS

### CellSP module discovery algorithm

CellSP identifies persistent subcellular spatial phenomena in the tissue by defining “gene-cell modules” as sets of genes with specific subcellular transcript spatial patterns across cells. This process is performed in two steps -

#### Subcellular pattern discovery

We utilize the existing tools InSTAnT [18] and SPRAWL [19] for subcellular pattern detection in individual cells. InSTAnT detects gene pair colocalization in single cells, employing the “Proximal Pairs” (PP) test to calculate colocalization p-values for each gene pair in every cell. It also assigns a global colocalization p-value to each gene pair, via the “Conditional Poisson Binomial” (CPB) test. We use a user-tunable threshold of 1*e*^−3^ on this p-value to limit the set of candidate gene pairs. A matrix *M*_*I*_ is constructed to store the PP test p-values with dimensions *n_cells_* × *n_genepairs_* where *n_genepairs_* is the number of candidate gene pairs. We then perform a negative log transformation on this matrix.

SPRAWL detects subcellular mRNA localization patterns and classifies them into four categories - peripheral, central, radial, and punctate. Each gene receives a statistical significance score (on a scale of −1 to 1) representing the strength of the spatial pattern in each cell. These scores are stored in four separate matrices *M_peripheral_*, *M_central_*, *M_radial_*, *M_punctate_* each with dimensions *n_cells_* × *n_genes_*.

#### Module discovery

Each entry of the matrices obtained from InSTAnT and SPRAWL represents the strength of pattern occurrence for each gene/gene-pair in each cell. To aggregate these patterns across genes and cells, we use the LAS [23] algorithm to perform biclustering on each of these matrices. The LAS algorithm iteratively searches for a submatrix (bicluster) with significantly high average value, removes the influence of selected submatrix from the original data matrix, and repeats the process until the desired number of submatrices are retrieved or no significant submatrices are found. The LAS significance score for a *k* × *l* submatrix *U* within a data matrix *X* of dimensions *m* × *n* is defined as

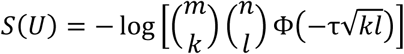

where Φ(⋅) denotes the cumulative distribution function (CDF) of the standard normal distribution, and τ denotes the average value of the submatrix *U*.

The reported submatrices are what we call “gene-cell modules” or “modules”. Due to the heuristic nature of the search, it is possible that a reported module is part of a larger module with a similarly high average value, but this larger module is not discovered. To mitigate this issue across reported modules, we implement a two-step module expansion process:

1. Cell Expansion – For each module, we iterate through all cells and add a cell to the module if the cell’s average value across the module’s genes exceeds the original module’s average value.
2. Module Mergers – Iterating over modules in descending order of significance score, we calculate the overlap coefficient for the genes of each module pair. The overlap coefficient is defined as

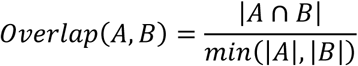

where *A* and *B* denote the gene sets of the two modules. If the overlap coefficient exceeds 0.667, then we consider merging the module pair by creating a new module that comprises the union of genes/gene pairs of the original pair and the union of cells of the original pair. If the new module’s significance score surpasses the module significance threshold, the merged module is retained, and the original pair of modules is removed. This process is repeated iteratively, prioritizing the merger of highly significant modules first, until no further mergers are possible. This merger step is useful to reduce the number of modules detected and combine modules of similar composition (high overlap in constituent genes) into one. This step is made optional.

The module discovery process (including module expansion) is conducted independently for modules derived from each of the five matrices *M*_*I*_, *M_peripheral_*, *M_central_*, *M_radial_*, *M_punctate_* that represent different kinds of subcellular patterns.

In the final reported list of gene-cell modules, two modules may comprise overlapping sets of cells and genes, allowing the same cells and genes to be part of multiple modules (**Fig. 1**). Modules detected from the SPRAWL-derived matrices *M_peripheral_*, *M_central_*, *M_radial_*, *M_punctate_* are reported as one group (their module identifiers have the suffix “_S” and a unique numeric prefix), while modules detected from the InSTAnT-derived matrix *M*_*I*_ are reported as another group (identifiers with suffix “_I” and a unique numeric prefix).

### Module Characterization

To investigate factors (other than cell types) that underlie or are associated with subcellular spatial phenomena, CellSP performs a two-level characterization of the detected modules using gene set enrichment analysis and predictive modeling.

Module Gene Characterization: The biological functions associated with module genes are characterized using PantherDB [74] for gene enrichment analysis. For each module, the module genes are used as the query gene set, while the gene panel of the ST assay serves as the background set. This analysis identifies pathways, biological processes, and molecular functions enriched in module genes.

Module Cell Characterization: To characterize the cells in each module, CellSP trains a classifier to distinguish between module cells (positive set) and all other cells (negative set). Each cell’s gene expression profile (total transcript count of each gene in that cell) is used as the cell’s feature vector, i.e., this is not a spatially resolved featurization. Since an ST assay of single-molecule resolution typically profiles a limited number of genes, the number of features is relatively modest. To address this, CellSP uses the Tangram tool [75] to impute the expression of additional genes in each cell of the ST data using scRNA-seq data of the same tissue, if available. The imputation process expands the gene set significantly, increasing the number of features from a few hundreds (in the original ST data) to several thousands (as available in scRNA-seq data). We construct an extended gene expression matrix by retaining the original expression values for genes present in the ST panel and incorporating the imputed expression values for genes absent from the panel. The extended gene expression matrix is used to train a Random Forest classifier of module cells. The top 20 most informative genes are identified based on their feature importance, which is quantified using SHAP [76] [77]. These genes, along with any other genes that exhibit a high Pearson correlation (r > 0.98) with them, are designated as “marker genes” of the module cells. These genes are then subjected to gene set characterization using PantherDB to identify pathways, biological processes, and molecular functions that characterize the module’s cells.

### Module Pattern Visualization

To help visualize modules defined by the five types of subcellular spatial patterns (four types identified by SPRAWL and the colocalization pattern identified by InSTAnT), we developed three complementary plotting techniques.

SPRAWL detects localization patterns (peripheral, central, radial or punctate) for each gene in each cell. To aggregate these spatial localization patterns across many cells, CellSP transforms each cell into a uniform representation within a unit circle. This transformation is achieved using the smallest enclosing circle algorithm to identify the smallest circle enclosing all transcripts in the cell. The circle is then centered at the origin and scaled to have a unit radius. For “central” and “peripheral” patterns, the unit circle is divided into ***C*** (default value of 5) concentric rings, and the proportion of module gene transcripts within each ring is calculated. The transcript abundance in a ring is then averaged across all cells of the module, and the ring is displayed with color shade depicting the average abundance. A similar visualization is constructed for all non-module cells and the two plots are placed side-by-side. For modules displaying “central” patterns, we expect higher abundance in the innermost rings, while “peripheral” patterns are anticipated to exhibit higher abundance in the outermost rings.

For modules defined by “punctate” or “radial” patterns, the unit circle representing an idealized view of a cell is divided into ***S*** (defaults to 10) sectors. Gene transcript density is computed in each sector for module genes. To account for the directional variability of these patterns, each cell’s idealized circle view is rotated such that the sector with the maximum density aligned with 0°. Following this alignment, densities are aggregated across cells by calculating mean densities for each sector. This results in a density distribution across sectors, with the highest density in the first sector by construction. This calculation is then repeated for non-module genes, and the density distributions (across sectors) for module genes and non-module genes are plotted overlaid on each other, in different colors. In this framework, module genes are expected to concentrate primarily in the first sector, while non-module genes are anticipated to exhibit more uniform distributions across sectors, despite having the highest density in the first sector.

InSTAnT detects colocalized gene pairs as those whose transcripts are in close proximity (within distance *d*) of each other. To visualize such patterns aggregated over cells, CellSP first constructs a *d*-radius neighborhood graph of transcripts for each module cell (where *d* equals the distance threshold parameter used in InSTAnT runs), calculates the number of neighboring transcript pairs for each pair of module genes and averages these counts across all module cells, thus obtaining a “proximity score” for that gene pair aggregating information across module cells. This process is then repeated over non-module cells to obtain a proximity score of the same gene pair but now aggregating information from non-module cells. The proximity enrichment score of the gene pair is then defined as the log ratio of proximity scores from module cells and non-module cells. High positive values of the proximity enrichment score indicate that the gene pair exhibits a greater degree of colocalization in module cells compared to non-module cells. To add further contrast, a randomly selected set of non-module (“control”) genes, equal in count to the module genes, is included in the visualization: a heatmap is constructed whose rows and columns represent module genes (top half of rows and left half of columns) and the selected control genes (bottom half of rows and right half of columns), and values depict proximity enrichment scores of gene pairs. The upper-left quadrant corresponds to pairs of module genes, the lower-right quadrant to pairs of control genes, and the remaining two quadrants correspond to a module gene paired with a non-module gene.

### Gene Set Enrichment Visualization

CellSP provides the ability to visualize the gene set enrichment reports generated by PantherDB using the Revigo tool [24]. Revigo summarizes lists of GO terms by clustering them based on their semantic similarity, identifying representative terms. These terms are visualized as circles in a scatterplot, where the circle size reflects the gene set size and the color intensity indicates statistical significance, with darker colors representing higher significance. This visualization highlights the importance, similarity, and uniqueness of terms, making it easier to interpret enrichment results.

We used a special type of Revigo plots to aggregate gene set enrichment reports across multiple detected modules (**Fig. 4d,e**; this is not a standard functionality of CellSP). We first collected all GO terms with a module association at a p-value below 0.01, across the modules, along with their p-values. This aggregated list of significant terms was then processed through Revigo, which calculated the principal component analysis (PCA) values for each term. Using these PCA coordinates, we generated a 2D histogram to visualize the aggregated GO terms across modules. The PCA values determined the positions of the bins in the plot and the number of modules each term was associated significantly with was represented by the color intensity of the corresponding bin. This visualization highlights the most commonly enriched biological themes across modules while preserving the semantic relationships among GO terms.

### Cross-Condition Module Comparison

#### Human Kidney Dataset

To compare the detected modules between conditions in the Human Kidney dataset, we assessed whether the subcellular patterns of modules from the cancer condition were present in the control tissue. We first performed min-max normalization of gene expression values for both the control and cancer datasets independently. For a gene-cell module detected in the cancer dataset, we calculated the total gene expression of module genes in each module cell and determined their minimum *P_min_* and maximum *P_max_* as percentiles of total gene expression in the population of all cells in the cancer dataset. Using these percentiles, we identified cells from the control dataset that have total gene expression (of module genes) between *P_min_* and *P_max_* percentile of the population of control cells. Thus, we identified a subset of the control cells whose total expression is similar to the module cells (which belong to the cancer data set). Subcellular patterns of module genes were then examined in these control cells and compared to the patterns in the module cells from the cancer dataset.

#### Mouse Alzheimer’s Disease Dataset

To identify changes in functionally similar (myelination-related) modules across conditions (AD model mice and WT mice), we asked whether two sets of modules (one set from each condition) differ in their gene composition. One way to answer this is based on how many modules in each condition include a specific gene, and whether the gene’s frequency of inclusion is different between conditions. This approach was used for **Fig. 5e**. A complementary approach is based on counting how many modules in each condition include a specific gene pair and comparing the proportion of such modules between conditions using a one-sided proportion test [78]. The test statistic for the proportion test is given by

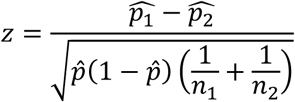

where 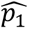 is the proportion for group 1, 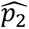 is the proportion for group 2, *p^* is the pooled proportion and *n*_1_, *n*_2_ are the sample sizes for group 1 and 2 respectively.

### Comparison with traditional analysis methods

To compare insights generated by our approach against traditional methods, we performed differential expression analysis and co-expression network analysis on the datasets. For differential expression analysis, we utilized Scanpy [79] to apply the Wilcoxon rank-sum test, comparing gene expression between the two conditions under investigation. This provided insights into genes with significantly altered expression levels across conditions.

For co-expression network analysis, we employed WGCNA [21] using the PyWGCNA framework [80]. Similar to our approach using CellSP, WGCNA was run independently for each cell cluster in the dataset. WGCNA assigns a module identity (class label) to each gene within a cluster based on its co-expression patterns. We used these module identities to calculate the frequency with which a gene pair is assigned to the same WGCNA module across the dataset. These frequencies were then compared between the conditions using a one-sided proportion test to assess condition-specific co-expression relationships.

### CellSP User Guide

CellSP provides flexible and tunable parameters that allow users to adapt analyses to their specific datasets and objectives. For InSTAnT, the distance threshold (***d***) is typically set to approximately 5% of the average cell diameter. The significance threshold (***α***) for the CPB p-values is set to 1*e*^−5^ by default. If this results in too few significant gene pairs (< 250), we recommend relaxing the threshold to 1*e*^−3^ or lower. To manage computational complexity, an additional parameter, ***K***, allows users to select the top ***K*** gene pairs for further analysis. For SPRAWL, we adapted the original scripts into CellSP to enable parallelization and use the default parameters. Similarly, LAS scripts from the implementation available in biclustlib [81] were parallelized and integrated into CellSP. LAS includes two primary adjustable parameters: ***N*** and ***RS***. The parameter ***N*** determines the number of modules to search and is set to a default value of 10. We recommend keeping this parameter between 5 to 50. The parameter ***RS*** specifies the number of randomized searches performed to enhance the quality of the biclusters. By default, ***RS*** is set to 50,000; however, for larger datasets (e.g., > 200 genes or > 10,000 cells), we recommend reducing this value to balance computational demands while maintaining reasonable bicluster quality. Visualization in CellSP is also highly customizable. Users can fine-tune the representation of subcellular patterns using parameters such as the number of concentric circles (***C***), the number of sectors (***S***), and the distance threshold (***d***). These settings provide flexibility in highlighting key spatial patterns. Details for the specific parameters used in each analysis can be found in **Supplementary Data 10**. On a dataset containing 10,000 cells and 248 genes, CellSP completes its analysis in approximately 3 hours. Additional runtime details can be found in **Supplementary Note 2** and **Supplementary Data 13.**

## Supporting information

Description of Supplementary Data

## DATA AVAILABILITY

The MERFISH dataset for the Mouse Preoptic Hypothalamus region [26] was obtained through direct communication with Dr. Jeffrey Moffitt. After excluding cells labeled with ambiguous cell types, the dataset comprises 5,149 cells distributed across 9 distinct cell types, with a gene panel of 135 genes.

The Xenium datasets are publicly available at 10x Genomics Datasets. We used the “Fresh Frozen Mouse Brain for Xenium Explorer Demo” [41], “Human Kidney Preview Data” [52], and the “Xenium In Situ Analysis of Alzheimer’s Disease Mouse Model Brain Coronal Sections from One Hemisphere Over a Time Course” [53] datasets for the mouse brain, human kidney, and Alzheimer’s disease experiments, respectively. The Mouse Brain dataset contains 162,033 cells grouped into 50 clusters based on the expression profiles of 248 genes. The Human Kidney dataset comprises 56,509 cells grouped into 19 clusters in the cancer tissue and 97,546 cells grouped into 21 clusters in the control tissue. The gene panel includes 377 genes. The Mouse Alzheimer’s dataset includes six tissue samples from two conditions—Wild Type and Alzheimer’s (TgCRND8 mouse model)—at three timepoints (2.5 months, 5.7 months, and 13+ months). The Alzheimer’s samples contain 53,908, 58,681, and 61,435 cells across the timepoints, while the Wild Type samples contain 58,230, 58,685, and 59,933 cells, with a gene panel of 347 genes.

## CODE AVAILABILITY

CellSP is open source and available at https://github.com/bhavaygg/CellSP.

## ACKNOWLEDGEMENTS

We thank Abhishek Ojha for his help in designing the statistical tests. This research utilized resources from PACE at the Georgia Institute of Technology, Atlanta, USA. Funding: This work was supported by the National Institutes of Health (R35GM131819 to S.S.), and Georgia Institute of Technology (Wallace H. Coulter Distinguished Faculty Chair: S.S.)

## Notes

### Competing Interest Statement

The authors have declared no competing interest.

### Summary of Updates

Revisions to the introduction sections to emphasize on limitations of current research and how our method builds on that.

