## Supplementary Figures for "CellSP: Module discovery and visualization for subcellular spatial transcriptomics data"

A

| Pattern | #Modules | Average #cells | Average #genes |
|---|---|---|---|
| Colocalization | 15 | 304 | 11 |
| Central | 5 | 132 | 9 |
| Peripheral | 3 | 167 | 12 |
| Punctate | 9 | 126 | 8 |
| Radial | 6 | 116 | 10 |

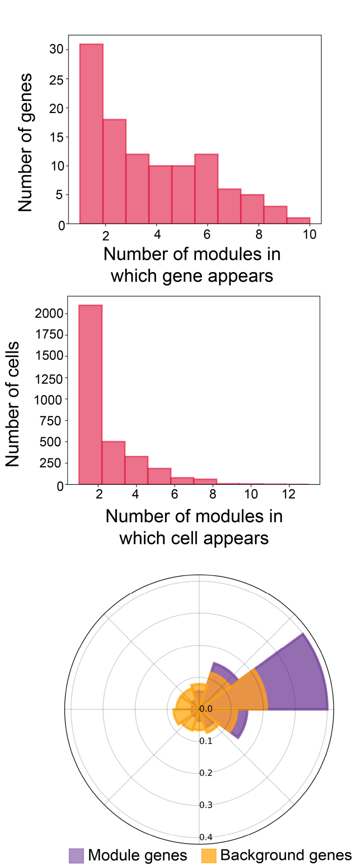


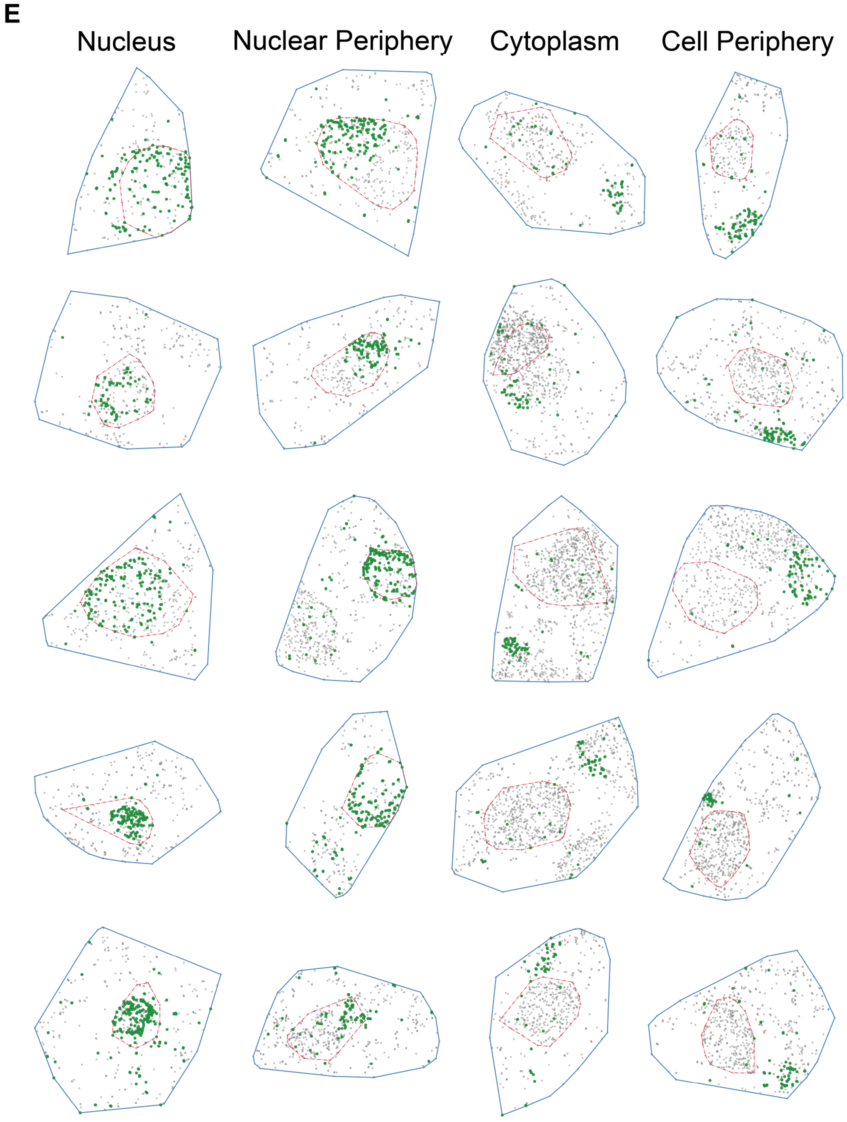


**Supplementary Figure 1**: **(A)** Summary table of the modules discovered by CellSP from MERFISH data on hypothalamic preoptic area in mouse brain. A module comprises of 9 genes and 138 cells on average. **(B)** Histogram showing the distribution of the number of modules in which a gene appears. A gene appears in less than four modules on average. **(C)** Histogram showing the distribution of the number of modules in which a cell appears. A cell appears in less than four modules on average. **(D)** CellSP visualization of the radial (or punctate) pattern for module "M11_S". The module genes (shown in purple) exhibit less spatial distribution, with a more concentrated localization towards one sector of the cell compared to non-module genes (shown in orange). **(E)** Direct visualization of example cells from the module "M0_I" which was discovered for exhibiting a colocalization pattern. The cells have been manually annotated with the subcellular localization of the modules genes. The colocalization pattern focuses on gene transcript proximity regardless of cellular location.

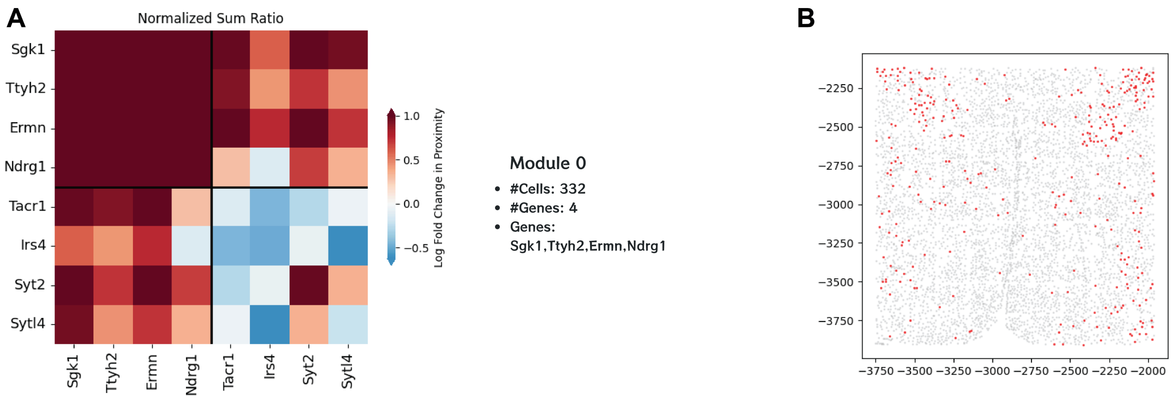


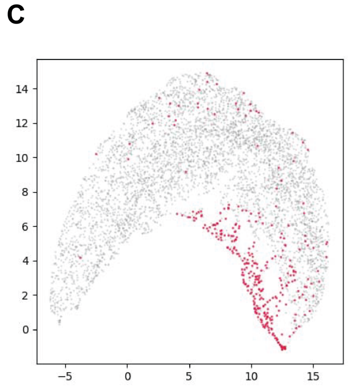


**D**

### GO Genes

| Term | P-Value |
|---|---|
| supramolecular fiber organization | 2.04e-02 |
| microtubule depolymerization | 2.84e-02 |
| sodium ion homeostasis | 2.84e-02 |
| microtubule polymerization or depolymerization | 2.84e-02 |
| negative regulation of microtubule polymerization or depolymerization | 2.84e-02 |

### GO Cells

| Term | P-Value |
|---|---|
| ensheathment of neurons | 1.40e-12 |
| axon ensheathment | 1.40e-12 |
| myelination | 1.32e-11 |
| regulation of glial cell differentiation | 3.52e-07 |
| regulation of oligodendrocyte differentiation | 4.17e-07 |

**Supplementary Figure 2:** CellSP creates a comprehensive webpage summary for all detected modules. We illustrate the report section for the module “M0_I” detected in the MERFISH data on hypothalamic preoptic area in mouse brain. The module was detected for exhibiting colocalization pattern, and the report showcases key details such as the identified subcellular pattern **(A)**, spatial distribution of module cells in the tissue **(B)**, UMAP visualization of the module cells **(C)**, and the functional characterization of the module **(D)**.

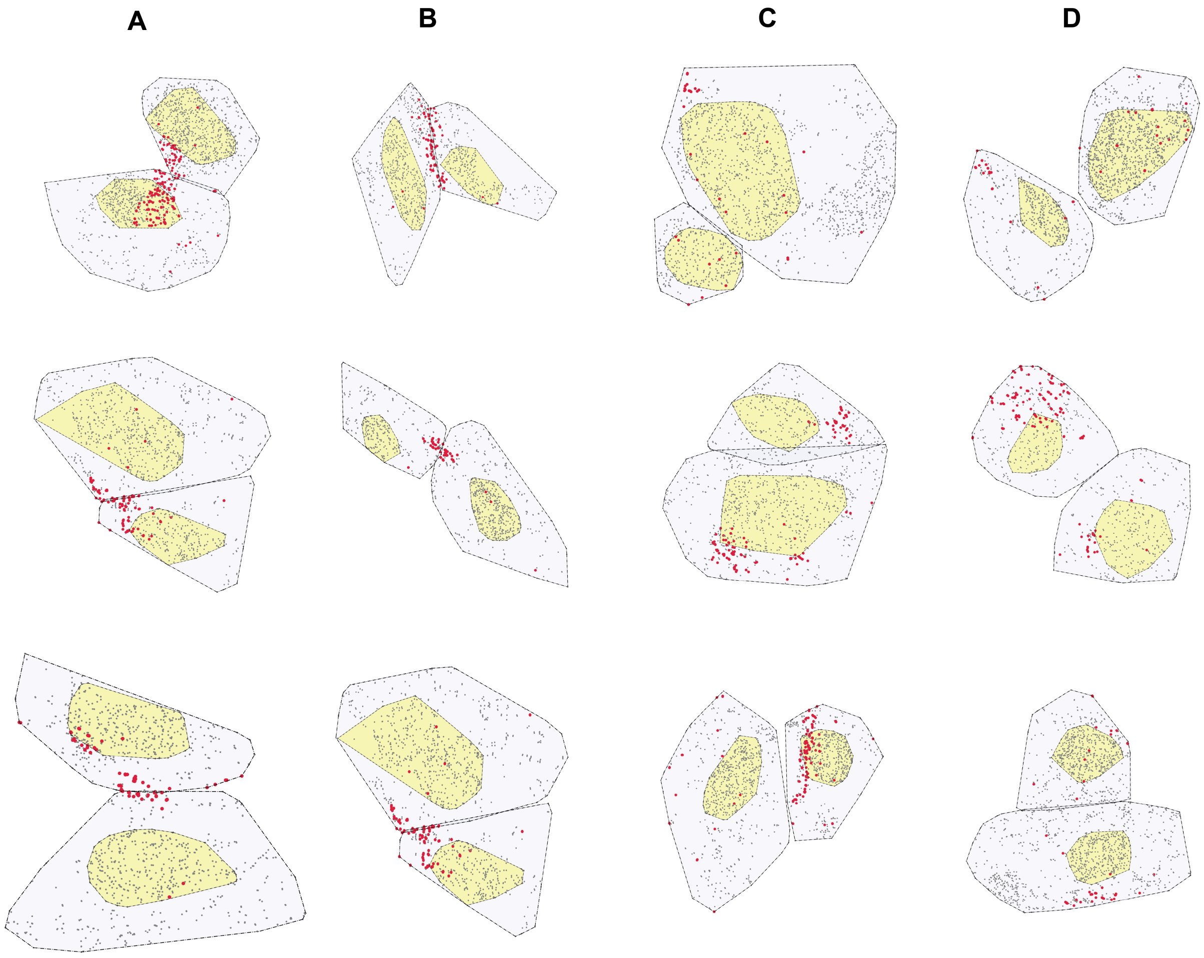


**Supplementary Figure 3:** CellSP discovered module "M0_S" in the MERFISH data on hypothalamic preoptic area is characterized by axonogenesis and cell-cell adhesion. To assess whether this characterization is reflected in the cellular localization of the genes and their spatial relationship with neighboring cells, we visualize module cells that are in close proximity and highlight the module genes. **(A, B)** Visualization of module cell pairs when they are **(A)** nearest neighbors and **(B)** second nearest neighbors. Module gene transcripts are highlighted in red. We observe that module genes localize predominantly to the cell contact boundaries, supporting their role in axonogenesis and cell-cell adhesion. **(C, D)** Visualization of randomly selected non-module cell pairs controlled for gene expression and cell type when they are **(C)** nearest neighbors and **(D)** second nearest neighbors. In contrast to the module cells, non-module gene transcripts do not exhibit similar subcellular localization patterns.

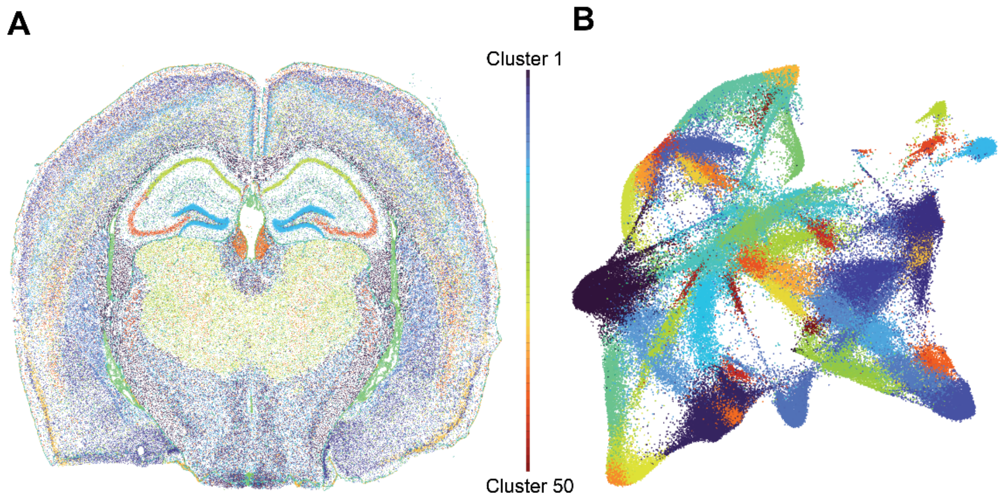


**Supplementary Figure 4: Overview of the Xenium whole mouse brain dataset. (A)** Spatial plot of all cells in the tissue colored by their cluster identity. **(B)** UMAP visualization of all cells colored by their cluster identity.

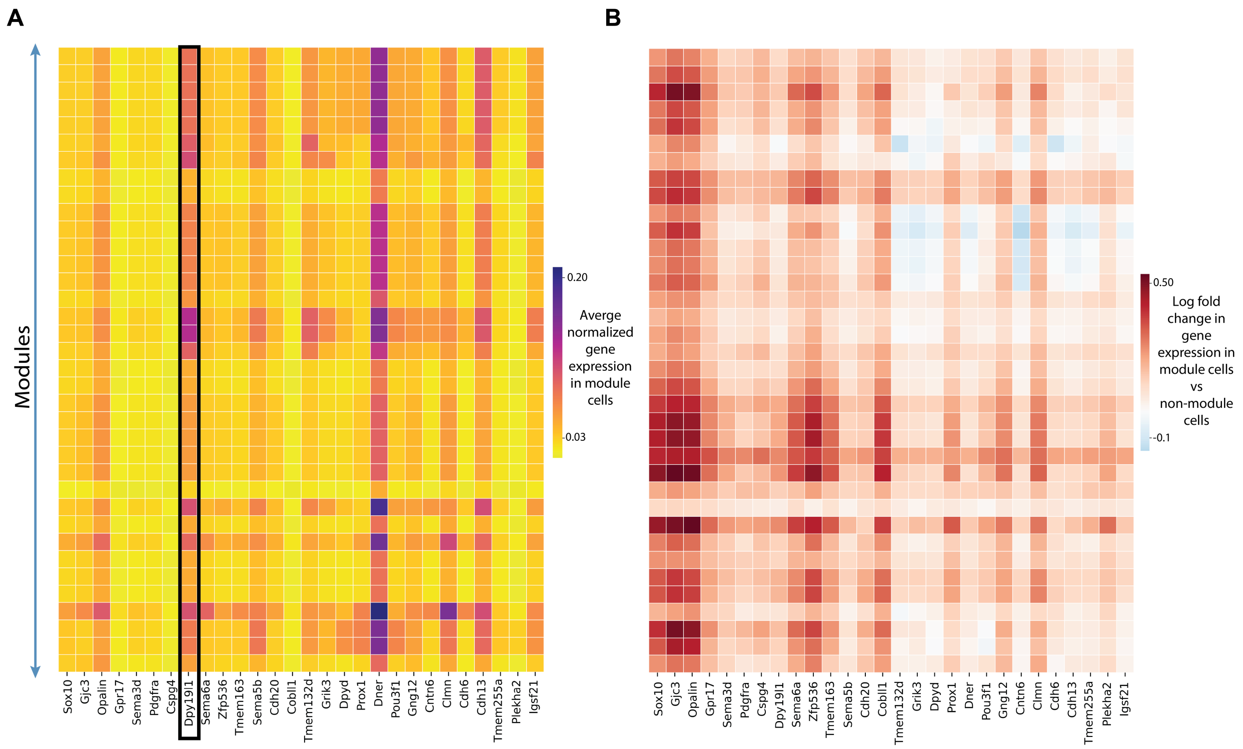


**Supplementary Figure 5: Comparing gene expression patterns in myelination-related modules from the Xenium whole mouse brain dataset.** We want to understand the influence of gene expression in the inclusion of the gene in a module. **(A)** Matrix showing the average normalized gene expression in module cells of each of the 36 myelination-related modules(rows) identified in the Xenium whole mouse brain dataset. We note that the gene expression, although indicative, is not solely predictive in a gene's inclusion in a module. For example, Dpy19l1 (dpy-19 like C-mannosyltransferase) (highlighted in black) shows high average expression in module cells, even though it is not part of the module genes. Conversely, it is also a part of the module genes when it shows lower average expression in the module cells. **(B)** Matrix showing the log fold change in gene expression in module cells versus non-module cells. The non-module cells are randomly chosen from the same expression cluster the module was detected in order to ensure expression similarity. We observe that module cells do tend to have a higher expression of module genes than non-module cells

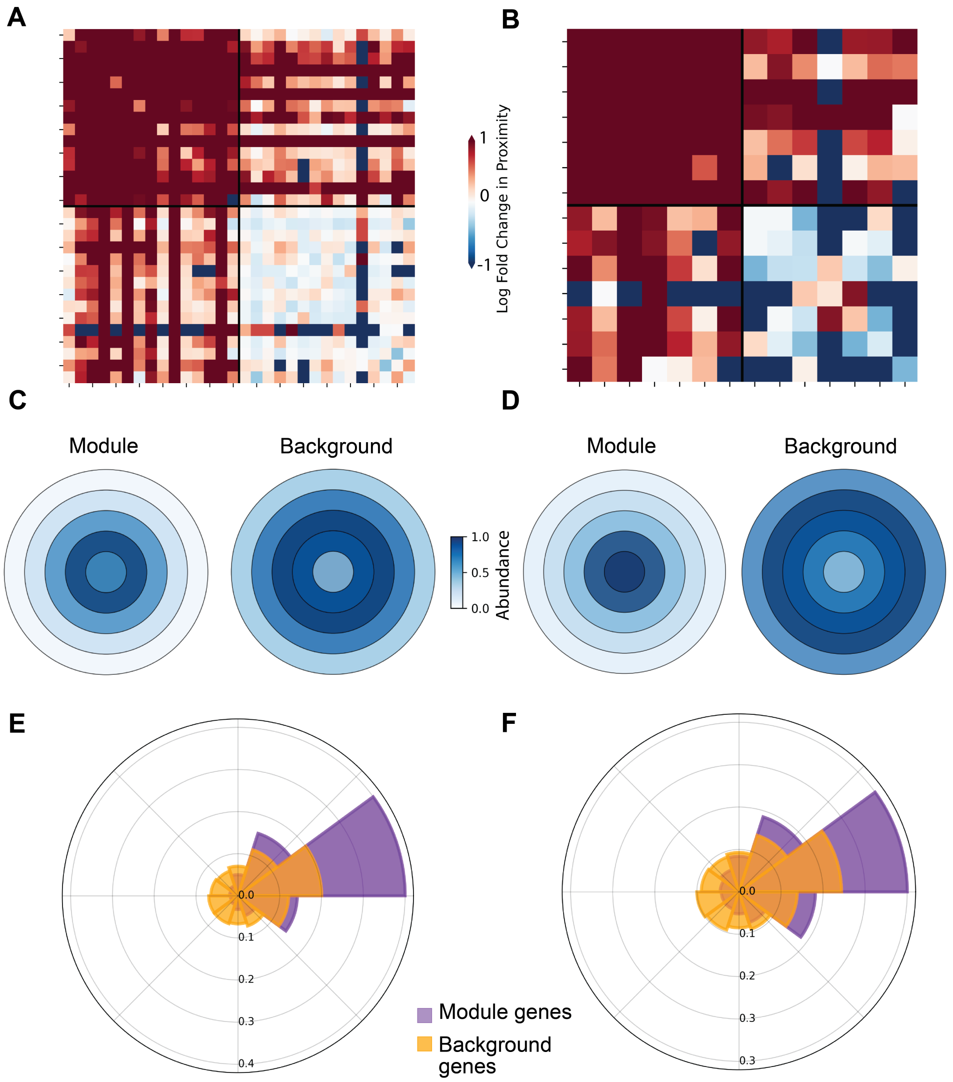


**Supplementary Figure 6: CellSP visualization of a subset of myelination-related modules discovered in the Xenium whole mouse brain dataset.** Colocalization patterns in modules “C3_M2_I” **(A)** and “C15_M2_I” **(B)** reveal that module genes are spatially clustered within module cells, exhibiting greater proximity compared to their distribution in non-module cells. Central localization patterns in modules “C3_M13_S” **(C)** and “C20_M12_S” **(D)** show that module genes tend to concentrate towards the center of module cells, contrasting with the more uniform distribution of non-module genes within the same cells. Radial and punctate patterns in modules “C3_M4_S” **(E)** and “C7_M4_S” **(F)** illustrate the preferential localization of module genes within specific radial sectors of module cells. (Note: In a module labelled “Cx_My_z” the prefix “Cx_” indicates the cluster number in which the module was detected.)

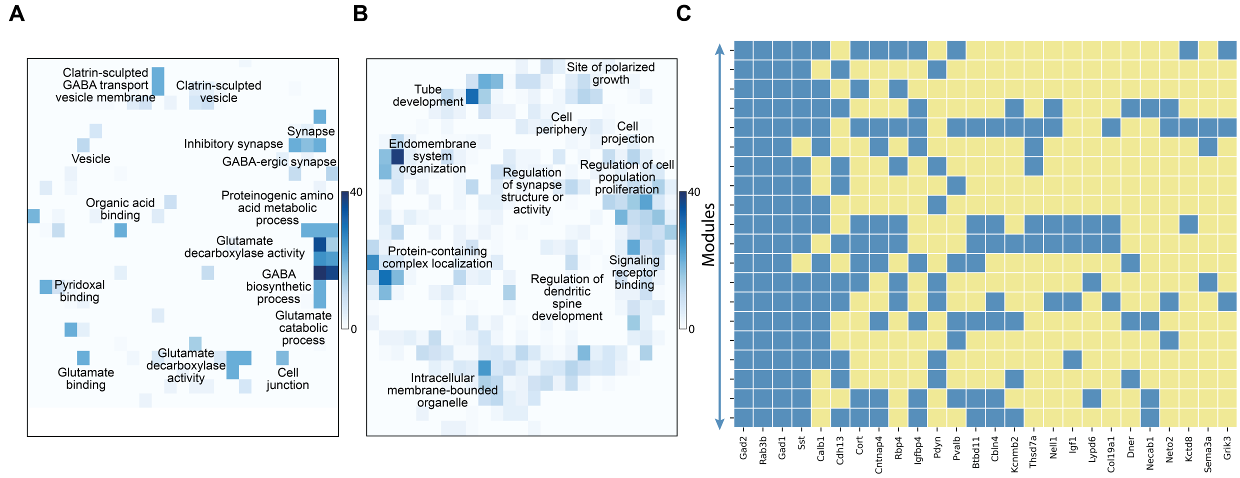


**Supplementary Figure 7: Gene enrichment and module composition analysis of the 20 GABA-ergic synapse-related modules discovered in the Xenium whole mouse brain dataset. (A)** REVIGO plot of biological processes associated with module genes aggregated over all 20 modules. Color intensity of a bin in this plot represents the number of significant module associations of one or more GO terms that map to that bin in the semantic space computed by REVIGO. **(B)** REVIGO plot of biological processes associated with gene markers of module cells aggregated over all 20 modules. **(C)** Matrix showing gene composition of each of the 20 modules; blue indicates a gene's inclusion in a module. Genes such as Gad1, Gad2, and Rab3d, frequently appear across modules, consistent with their critical roles in regulating GABA signaling.

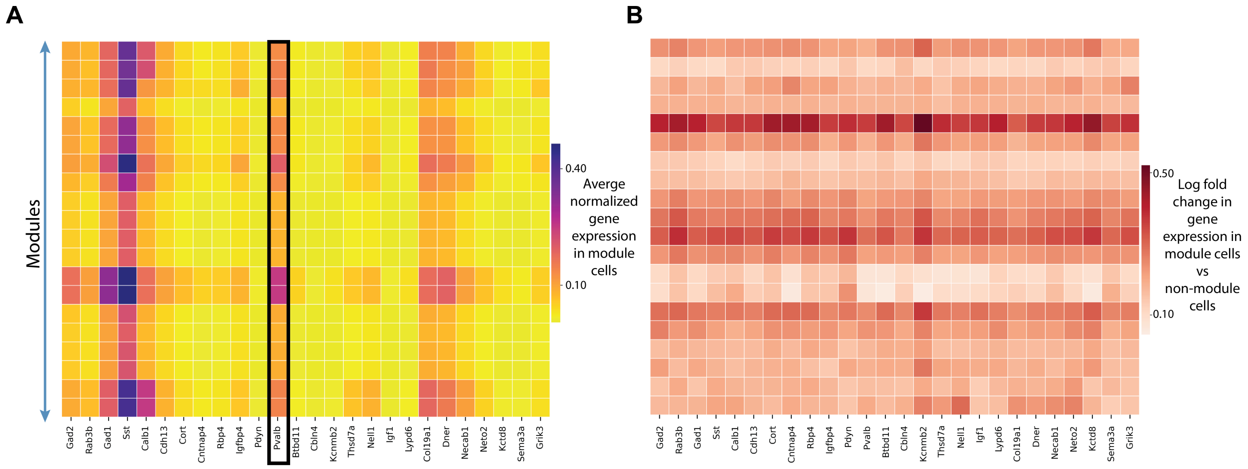


**Supplementary Figure 8: Comparing gene expression patterns in GABA-ergic synapse-related modules from the Xenium whole mouse brain dataset. (A)** Matrix showing the average normalized gene expression in module cells of each of the 20 GABA-ergic synapse-related modules (rows). Similar to Dpy19l1 in the myelination-related modules, we see variations in the expression genes like Pvalb (parvalbumin) whose gene-expression trends are not indicative of its inclusion in the module. **(B)** Matrix showing the log fold change in gene expression in module cells versus non-module cells. The non-module cells are randomly chosen from the same expression cluster the module was detected in order to ensure expression similarity. Similar to the myelination-related modules (**Figure S5B**), we see that module genes tend to have higher expression in module cells compared to non-module cells.

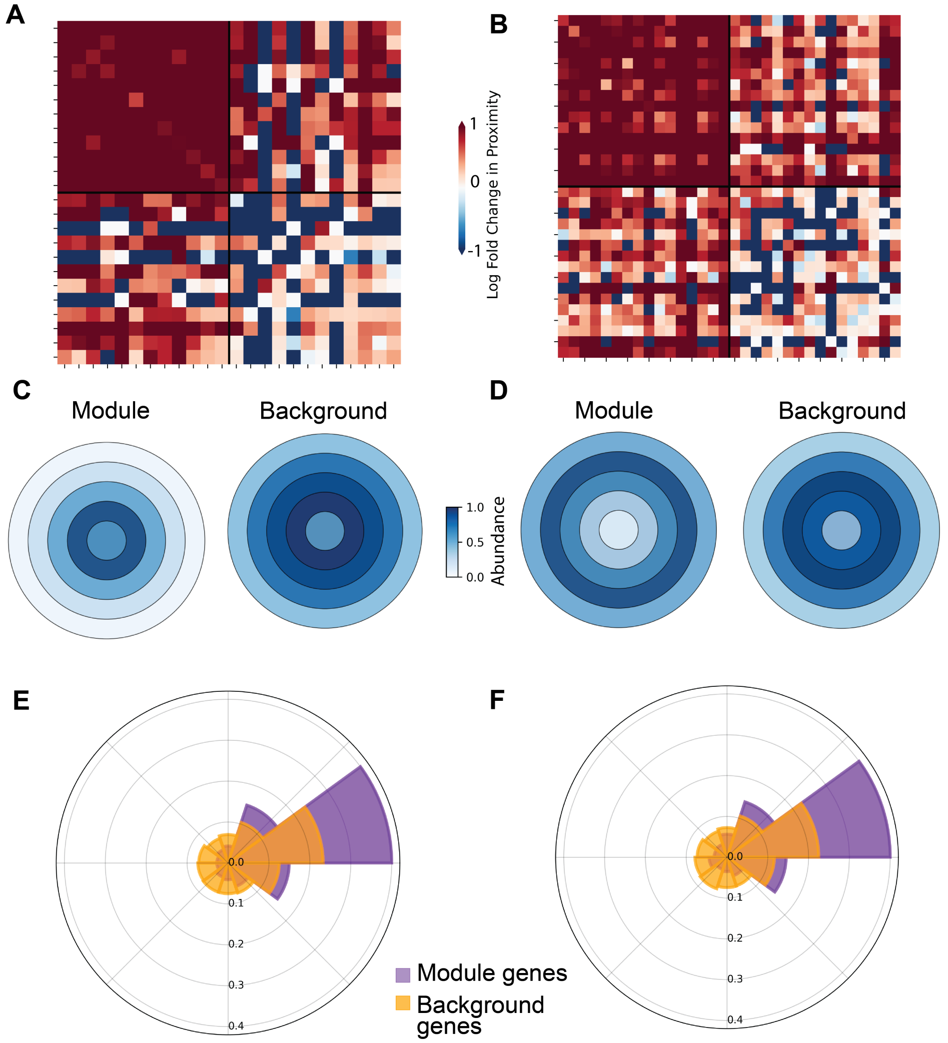


**Supplementary Figure 9: CellSP visualization of a subset of GABA-ergic synapse-related modules discovered in the Xenium whole mouse brain dataset.** Colocalization patterns in modules “C3_M8_I” **(A)** and “C6_M1_I” **(B)** demonstrate that module genes are more spatially clustered within module cells, exhibiting greater proximity compared to their distribution in non-module cells. Central and peripheral localization patterns in modules “C7_M10_S” **(C)** and “C42_M0_S” **(D)** reveal that module genes in “C7_M10_S” are concentrated towards the cell center, while those in “C42_M0_S” are localized at the cell periphery. In contrast, non-module genes show a more uniform distribution in the same cells. Radial and punctate patterns in modules “C15_M5_S” **(E)** and “C15_M8_S” **(F)** highlight the spatial organization of module genes into specific radial sectors within module cells.

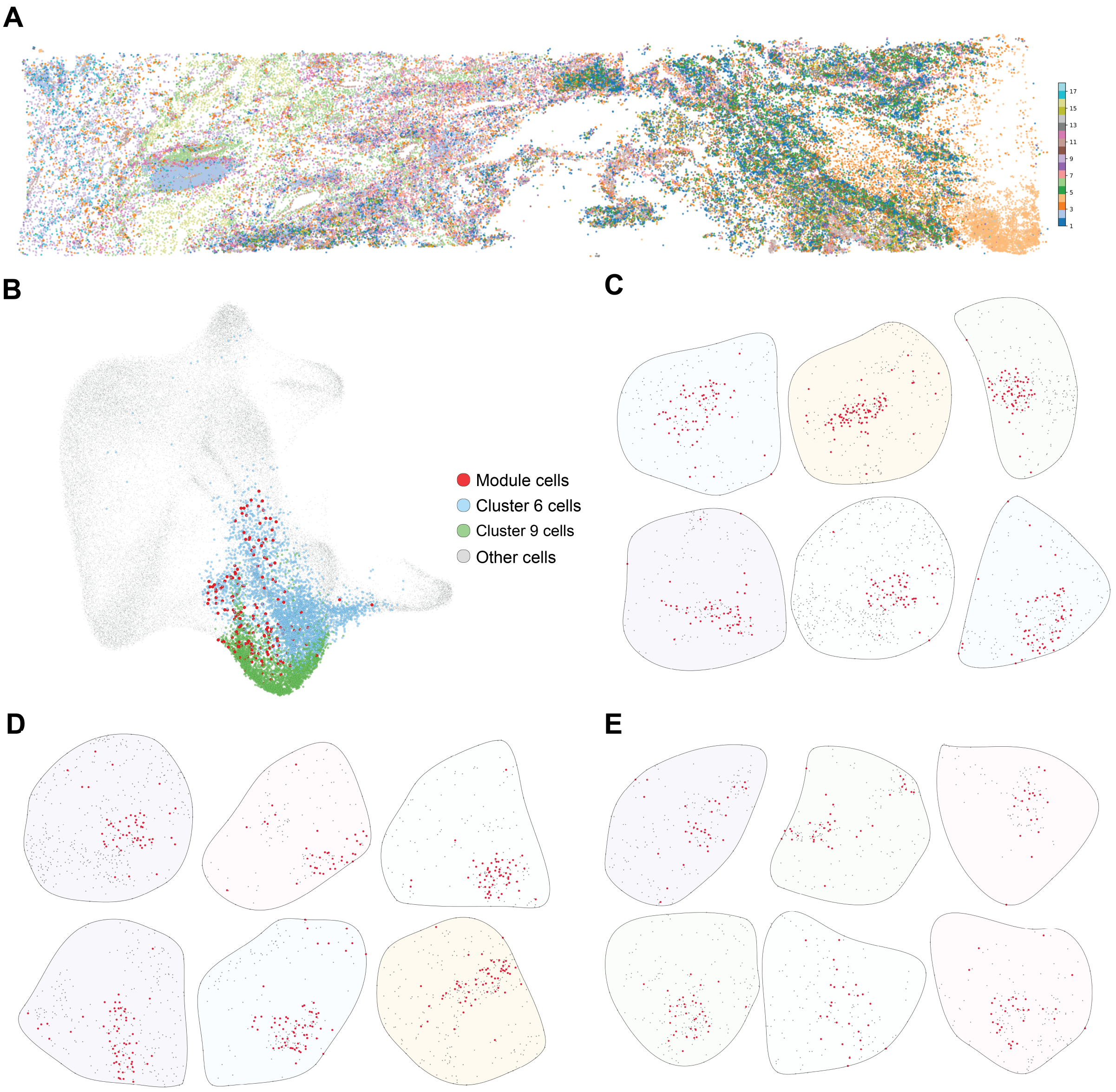


**Supplementary Figure 10: (A)** Spatial plot of all cells in the kidney PRCC tissue (Xenium data) colored by the cluster identity. **(B)** UMAP plot of all cells in the kidney PRCC tissue (Xenium data) highlighting cells from expression cluster 6 in blue, cells from expression cluster 9 in green and cells of any of the 6 CellSP-detected immune-response related gene-cell modules in red. **(C-E)**. Direct visualization of a subset of cells from modules "C6_M5_S," "C6_M0_S," and "C6_M3_I," respectively, showcasing the distinct subcellular patterns captured. Module genes, shown in red, exhibit a central concentration in "C6_M5_S" **(C)**, a radial distribution in "C6_M0_S" **(D)**, and spatial proximity in "C6_M3_I" **(E)**.

| Mouse | #clusters | #modules | #modules per cluster |
|---|---|---|---|
| WT 2.5 | 20 | 244 | 12 |
| WT 5.7 | 18 | 231 | 12 |
| WT 13.5 | 25 | 279 | 11 |
| Alzheimer's 2.5 | 21 | 235 | 11 |
| Alzheimer's 5.7 | 29 | 266 | 9 |
| Alzheimer's 17.9 | 20 | 212 | 10 |

**Supplementary Figure 11: Summary table of the modules detected by CellSP in each condition and timepoint of the Xenium Alzheimer's dataset.** We do not observe any variation in the number of modules detected between wildtype (WT) vs Alzheimer's (AD) mice.

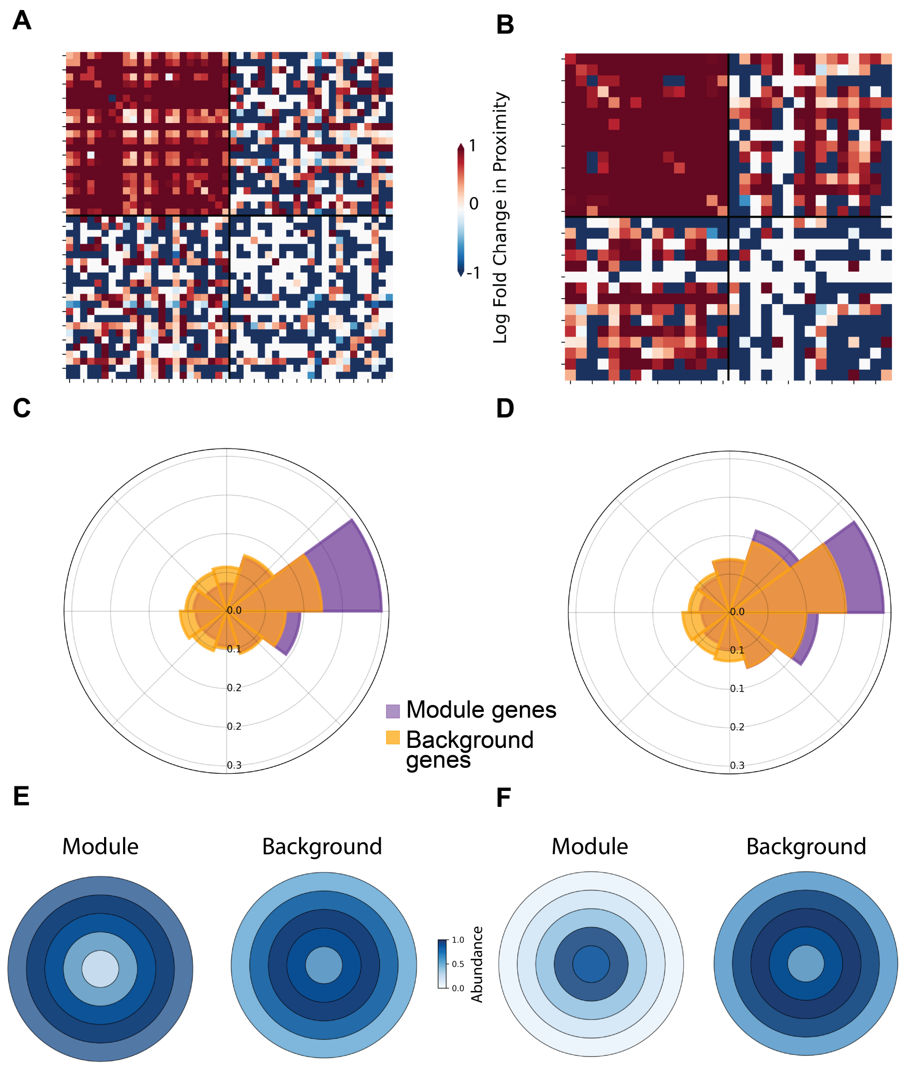


**Supplementary Figure 12: CellSP visualization of a subset of Picalm containing modules in the TgCRND8 mice.** Colocalization patterns in modules “C3_M1_I” **(A)** at timepoint 1 and “C3_M1_I” **(B)** at timepoint 2 demonstrate greater spatial proximity of module genes within module cells compared to their distribution in non-module cells. Radial and punctate patterns in modules “C13_M1_S” **(C)** at timepoint 1 and “C9_M6_S” **(D)** at timepoint 2 illustrate the organization of module genes into specific radial sectors within module cells. Peripheral and central localization patterns in modules “C22_M0_S” **(E)** at timepoint 1 and “C14_M7_S” **(F)** at timepoint 2 show that module genes in “C22_M0_S” localize towards the cell periphery, while those in “C14_M7_S” concentrate at the cell center. In both cases, non-module genes display a more uniform distribution within the same cells.

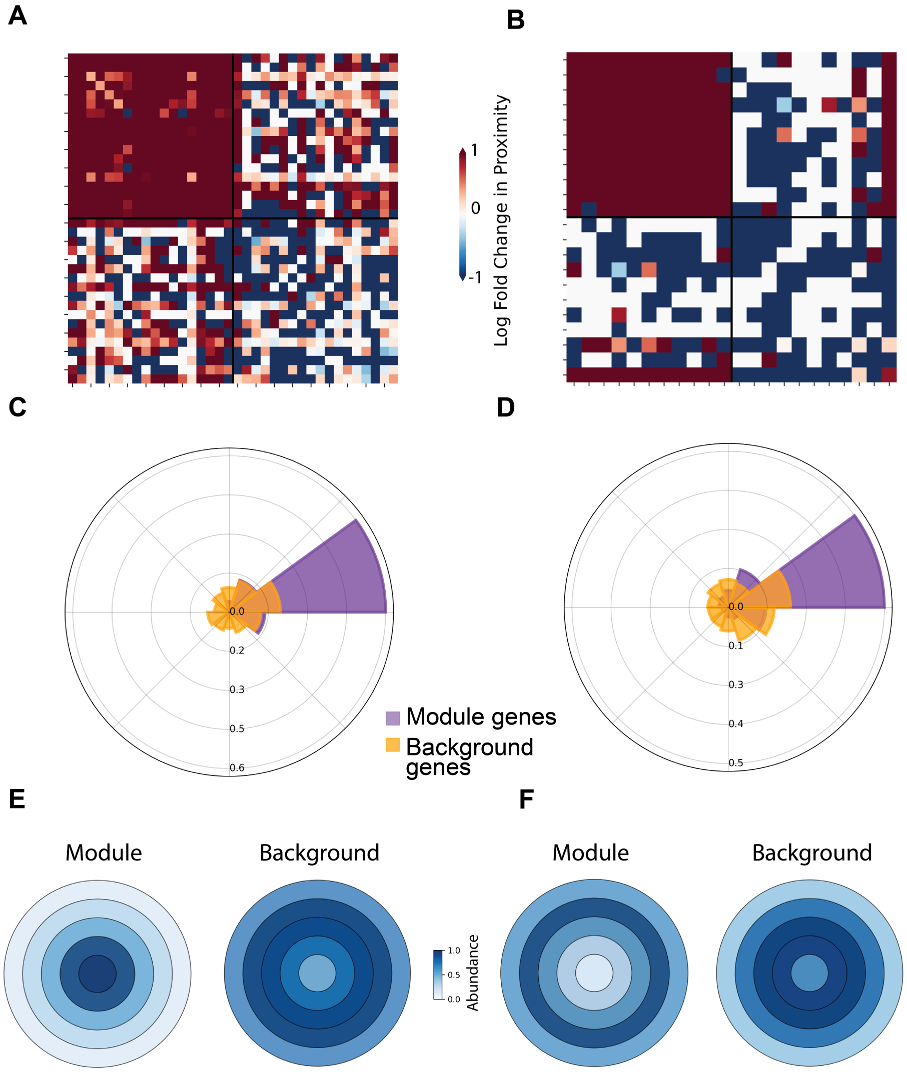


**Supplementary Figure 13: CellSP visualization of a subset of Trem2 containing modules in the TgCRND8 mice.** Colocalization patterns in modules "C1_M2_I" **(A)** and "C9_M1_I" **(B)** at timepoint 3 demonstrate greater spatial proximity of module genes within module cells compared to their distribution in non-module cells. Radial and punctate patterns in modules "C1_M2_S" **(C)** and "C16_M7_S" **(D)** at timepoint 3 show the concentration of module genes in specific radial sectors within the module cells. Central and peripheral localization patterns in modules "C3_M12_S" **(E)** and "C16_M0_S" **(F)** at timepoint 3 show that module genes in "C3_M12_S" are concentrated towards the cell center, while those in "C16_M0_S" are localized at the cell periphery. Non-module genes, in contrast, exhibit a more uniform distribution within the same cells.

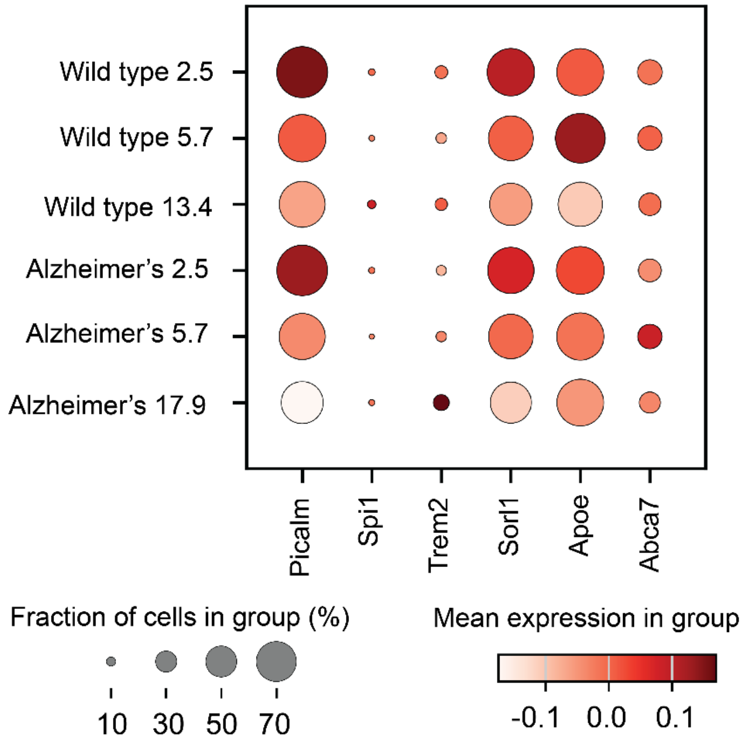


**Supplementary Figure 14: Analyzing the influence of gene expression on module detection in the Xenium Alzheimer's dataset.** Differential expression of AD risk genes across the different timepoints and conditions is visualized in a dot plot. The size of the circle represents the fraction of cells in the group that express the gene while the color represents the mean expression in the group. Consistent with our previous observations, we note that module detection by CellSP is influenced by, but not entirely dependent on, gene expression levels. For example, Trem2 shows higher expression in AD at timepoint 3, aligning with increased module detection. In contrast, Picalm has similar expression levels in both WT and AD at timepoint 1 but is detected in significantly greater number of modules in AD.
