## Supplementary Notes for "CellSP: Module discovery and visualization for subcellular spatial transcriptomics data"

**Supplementary Note 1: Comparison of CellSP with existing module discovery approaches.** The authors of InSTAnT presented two distinct module discovery approaches to identify modules of co-localizing gene pairs. The first approach, called Global Colocalization Clustering (GCC), uses hierarchical clustering to identify a set of genes enriched in pairwise colocalization (across cells). However, the gene-gene co-localization phenomena represented by a GCC-reported module may not manifest in the same cells, making such modules fundamentally different from CellSP's gene-cell modules. The second approach, Frequent Subgraph Mining (FSM), focuses on identifying sets of cells in which a specific number of genes exhibit mutual colocalization, revealing more localized patterns of gene interaction. This definition of modules is closer to CellSP's notion of modules, but is considerably more restrictive and less rigorous statistically compared to the latter, as we show below.

We compared both GCC and FSM to CellSP, applying all three approaches to the MERFISH dataset of the hypothalamic preoptic area (POA). CellSP identified 14 biclusters (gene-cell modules) encompassing 98 unique genes (out of the panel of 135 genes). On average, each module contained approximately 11 genes and was detected in 304 cells. CellSP also provides functional characterization of the module genes as well as the gene markers of the module cells. In contrast, GCC reported 10 clusters (**Data S11**); however, the results were dominated by a single large cluster containing 118 genes (out of 135 genes), making it challenging to draw biologically meaningful conclusions. This outcome suggests that GCC's reliance on global colocalization trends may limit its effectiveness in tissues with highly heterogeneous gene expression, such as the POA. GCC may instead be better suited to datasets derived from cell lines or tissues with more uniform gene expression profiles.

We used FSM to report modules consisting of four genes ($\boldsymbol{n}$), with the number of edges ($\boldsymbol{nE}$) in the subgraphs ranging from 4 to 6. (These two numbers are required parameters for using FSM.) For these configurations, FSM identified 1717, 351, and 38 modules, respectively (**Data S12**). The corresponding numbers of unique genes in these modules were 61, 39, and 19, respectively. In practical terms, the settings $\boldsymbol{nE}$ = 4,5 produce an unmanageably large number of modules (over 350) for a biologist to sift through. The setting $\boldsymbol{nE}$ = 6 produces a reasonable output (38 modules) but these modules are highly redundant (any gene occurs in two modules on average). Moreover, the discovered modules (at $\boldsymbol{nE}$ = 6) spanned relatively few cells, with only one reasonably size module (72 cells) and the rest being < 30 cells (13 cells on average) in size. Thus, the FSM method of module discovery is highly restrictive in what it considers to be a module, reducing our ability to recover biologically meaningful spatial patterns in the data set. It also lacks the statistical grounding of CellSP. The high redundancy among reported modules is also a practical problem, as it requires users to manually aggregate related modules, creating significant overhead. Furthermore, FSM necessitates multiple runs to explore different module sizes and levels of connectedness, making it a labor-intensive and time-consuming method to use effectively.

In contrast, CellSP leverages biclustering to directly identify co-localizing gene modules without the need for extensive parameter tuning or post-processing. This makes CellSP a more reliable and scalable approach, particularly for analyzing complex spatial transcriptomic datasets. By efficiently balancing interpretability and computational feasibility, CellSP provides an advantageous framework for discovering biologically meaningful gene modules.

**Supplementary Note 2: Scalability of CellSP.** To evaluate the scalability of CellSP with increasing tissue size, we performed analyses on subsets of cells sampled from the Xenium Mouse Brain dataset (**Data S13**). For these runs, the parameters were set as follows: $\boldsymbol{d}$ = 2, $\boldsymbol{\alpha}$ = 1e-5, $\boldsymbol{K}$ = 250, $\boldsymbol{N}$ = 10 and $\boldsymbol{RS}$ = 5000.

As the number of cells increased, we observed that a significant proportion of computational time was consumed during cell characterization, a step involving the training of a random forest model to differentiate between module and non-module cells. To address the class imbalance between module and non-module cells, we sampled non-module cells in each iteration and identified the most important features from the trained models. However, this iterative process became a computational bottleneck as the dataset size grew. To improve runtime efficiency, we suggest reducing the number of iterations in this step, which strikes a balance between computational cost and accurate feature selection.
