## Supplementary material for "CellSP: Module discovery and visualization for subcellular spatial transcriptomics data": Description of Supplementary Data

**Data S1:** Modules detected by CellSP in the preoptic area (POA) of the hypothalamus in the mouse brain. The "Module_ID" column provides a unique identifier for each module in the format "Mx_y," where x denotes the module number and y represents the tool used to identify the module (InSTAnT or SPRAWL). The "Pattern" column specifies the subcellular pattern displayed by the module. The "Genes" column lists the genes in each module, while "#genes" indicates the number of genes in the module. Similarly, the "uIDs" column lists the unique identifiers of the cells in each module, and "#cells" shows the number of cells in the module. The "GO Module" column reports the most significant Gene Ontology (GO) term associated with the module genes, and the "GO Cell" column highlights the most significant GO term associated with the marker genes of the module cells. The "LAS score" column reflects the significance score assigned to the module by the LAS algorithm. Related to Figure 2 and Figure 3

**Data S2:** CellSP detected modules in the Xenium whole mouse brain dataset. The semantics of all columns remain unchanged from Data S1 except for "Module ID". In this dataset, the "Module ID" column includes an additional prefix "Cz_", where z denotes the expression cluster to which the module belongs. related to Figure 4

**Data S3:** CellSP detected myelination related modules in the Xenium whole mouse brain dataset. Columns have the same semantics as in Data S2. Related to Figure 4

**Data S4:** CellSP detected GABA-ergic synapse related modules in the Xenium whole mouse brain dataset. Columns have the same semantics as in Data S2. Related to Figure 4

**Data S5:** CellSP detected modules in the human kidney PRCC cancer sample. Columns have the same semantics as in Data S2. Related to Figure 5

**Data S6:** CellSP detected modules in the human kidney control sample. Columns have the same semantics as in Data S2. Related to Figure 5

**Data S7:** CellSP detected modules in the mouse brain Alzheimer's samples. Columns have the same semantics as in Data S2 with the addition of the column, "Mouse", which denotes the condition and timepoint of the tissue in which the module was detected. Related to Figure 5

**Data S8**: Proportion test results comparing the co-occurrence of genes in a module between control and Alzheimer's samples. The columns "G1" and "G2" represent the gene pairs whose module co-occurrence is being analyzed across the two conditions. The columns "x_ctrl" and "x_alz" indicate the number of modules where "G1" and "G2" co-occur in control and Alzheimer's samples, respectively. The columns "n_ctrl" and "n_alz" specify the number of modules associated with the Gene Ontology (GO) term "ensheathment of neurons" in control and Alzheimer's samples. The columns "pval_ctrl" and "zscore_ctrl" report the p-value and z-score from the one-sided proportion test for greater co-occurrence in control tissues. Similarly, "pval_alz" and "zscore_alz" provide the p-value and z-score from the one-sided proportion test for greater co-occurrence in Alzheimer's tissues. Related to Figure 5

**Data S9:** Proportion test results comparing module co-occurrence of Olig2 with its group specific partner genes across each timepoint of the control and Alzheimer's samples. The columns retain the same semantics as in Data S8, with the addition of "Timepoint", which specifies the timepoint of the tissues being compared. P-values less than 0.05 have been highlighted. Related to Figure 5

**Data S10:** CellSP parameters used for the datasets used. "Dataset" refers to the dataset. Columns "d", "α", "K", "N" and "RS" denote the CellSP parameters. Related to Methods.

**Data S11:** Modules detected by the GCC method described in InSTAnT. "Cluster" refers to the cluster identity assigned to the gene module by GCC. Columns "Genes" and "#Genes" denote the genes in the module and the number of genes in the module, respectively. Related to Note S1

**Data S12:** Modules detected by the FSM method described in InSTAnT. "Support" refers to the number of cells in which the gene module was detected. "Vertices" denotes the genes in the module. "Number of Edges" indicates how many of the genes are colocalized in the "Support". Related to Note S1.

**Data S13:** Benchmarking scalability of CellSP. Columns “#cells” and ”#genes” denote the number of cells and the number of genes that are randomly sampled from the Xenium whole mouse brain dataset. “Total Time” denotes the total runtime of CellSP. “InSTAnT Time” and “SPRAWL Time” denote the time taken by InSTAnT and SPRAWL respectively. “Biclustering Time” denotes the time taken to run biclustering on all five matrices produced by InSTAnT and SPRAWL. “Modelling Time” denotes the time taken to train a Random Forest model to differential module cells from non-module cells and extract genes that mark the module cells. “GO Enrichment Time” denotes the time take to run GO enrichment analysis using PantherDB. Related to Note S2
